# Signature morpho-electric properties of diverse GABAergic interneurons in the human neocortex

**DOI:** 10.1101/2022.11.08.515739

**Authors:** Brian Lee, Rachel Dalley, Jeremy A Miller, Thomas Chartrand, Jennie Close, Rusty Mann, Alice Mukora, Lindsay Ng, Lauren Alfiler, Katherine Baker, Darren Bertagnolli, Krissy Brouner, Tamara Casper, Eva Csajbok, Nick Dee, Nicholas Donadio, Stan L.W. Driessens, Tom Egdorf, Rachel Enstrom, Anna A Galakhova, Amanda Gary, Emily Gelfand, Jeff Goldy, Kristen Hadley, Tim S. Heistek, Dijon Hill, Nelson Johansen, Nik Jorstad, Lisa Kim, Agnes Katalin Kocsis, Lauren Kruse, Michael Kunst, Gabriela Leon, Brian Long, Matthew Mallory, Michelle Maxwell, Medea McGraw, Delissa McMillen, Erica J Melief, Gabor Molnar, Marty T Mortrud, Dakota Newman, Julie Nyhus, Ximena Opitz-Araya, Trangthanh Pham, Alice Pom, Lydia Potekhina, Ram Rajanbabu, Augustin Ruiz, Susan M Sunkin, Ildiko Szots, Naz Taskin, Bargavi Thyagarajan, Michael Tieu, Jessica Trinh, Sara Vargas, David Vumbaco, Femke Waleboer, Natalie Weed, Grace Williams, Julia Wilson, Shenqin Yao, Thomas Zhou, Pal Barzo, Trygve Bakken, Charles Cobbs, Richard G. Ellenbogen, Luke Esposito, Manuel Ferreira, Nathan W Gouwens, Benjamin Grannan, Ryder P. Gwinn, Jason S. Hauptman, Rebecca Hodge, Tim Jarsky, C.Dirk Keene, Andrew L. Ko, Boaz Levi, Jeffrey G. Ojemann, Anoop Patel, Jacob Ruzevick, Daniel L. Silbergeld, Kim Smith, Jack Waters, Hongkui Zeng, Jim Berg, Natalia A. Goriounova, Brian Kalmbach, Christiaan P.J. de Kock, Huib D Mansvelder, Staci A Sorensen, Gabor Tamas, Ed S. Lein, Jonathan T Ting

## Abstract

Human cortical interneurons have been challenging to study due to high diversity and lack of mature brain tissue platforms and genetic targeting tools. We employed rapid GABAergic neuron viral labeling plus unbiased Patch-seq sampling in brain slices to define the signature morpho-electric properties of GABAergic neurons in the human neocortex. Viral targeting greatly facilitated sampling of the SST subclass, including primate specialized double bouquet cells which mapped to two SST transcriptomic types. Multimodal analysis uncovered an SST neuron type with properties inconsistent with original subclass assignment; we instead propose reclassification into PVALB subclass. Our findings provide novel insights about functional properties of human cortical GABAergic neuron subclasses and types and highlight the essential role of multimodal annotation for refinement of emerging transcriptomic cell type taxonomies.

**One Sentence Summary:** Viral genetic labeling of GABAergic neurons in human *ex vivo* brain slices paired with Patch-seq recording yields an in-depth functional annotation of human cortical interneuron subclasses and types and highlights the essential role of multimodal functional annotation for refinement of emerging transcriptomic cell type taxonomies.

## Main Text

Recent work on the human brain has shed new light on transcriptomic cell type diversity, including the identification of novel cell types and finer distinctions than previously recognized (*1*–*11*). Single cell transcriptomics has taken center stage as the leading scalable technology for understanding cell type diversity and is expected to soon yield a comprehensive parts list and unifying reference for the human brain. While having a complete parts list bounds the problem of cell type diversity, a deeper understanding of the phenotypic properties of transcriptomic cell types is necessary to provide mechanistic insights into their role in cognitive function and disease. Although it is hypothesized in transcriptomic cell type classification that differentially expressed genes within each transcriptomic type (t-type) underlie distinct morphological and functional properties, direct measurements are lagging well behind the pace of transcriptomic characterization. Additionally, there exists unresolved ambiguity across, and diversity within, transcriptomic types that remains underexplored for many different brain regions and species.

The Patch-seq method (patch-clamp electrophysiology plus single cell RNA-sequencing) provides the most direct means to determine the fundamental morpho-electric properties of transcriptomic neuronal types, evaluate distinctness between types and explore gene-function relationships (*12*–*14*). Already this approach has been used successfully to characterize the most abundant glutamatergic pyramidal neuron types in the human supragranular cortex (*15*). This work provided deeper functional context regarding the greater diversity of human glutamatergic pyramidal neuron types relative to mouse supragranular cortex as well as elucidation of the transcriptomic identity of human cortical cell types with selective vulnerability in disease. In addition, gradients in gene expression were largely mirrored by gradients in other measured morpho-electric features, consistent with other recent Patch-seq studies of mouse cortical neuron types (*16*, *17*). It has also been possible to apply Patch-seq to measure the multimodal cellular properties of cortical layer 5 extratelencephalic projecting (L5 ET) neurons in the human cortex (*2*, *18*), and to apply homology mapping based on analysis of single cell transcriptomes for comparing the cellular features of this subcortically projecting cortical neuron type across mammalian species. Characterization of these relatively sparse human L5 ET neurons in Patch-seq experiments was achieved by examining soma size, shape, and laminar position, thus making systematic targeting tractable.

GABAergic neurons are crucial for modulating and tuning neuronal circuits (*19*) and their dysfunction is at the forefront of a variety of human brain disorders (*20*–*22*), yet a comprehensive characterization of human cortical GABAergic neuron types has proven technically far more challenging. This stands in contrast to decades of impressive advances in the mouse model, where development and refinement of a multitude of viral and transgenic targeting and *in vivo* perturbation strategies have facilitated rapid progress (*23*). GABAergic neurons in the cortex historically have been subdivided into four major subclasses based on the neuro-chemical markers they express: parvalbumin (PVALB), somatostatin (SST), vasoactive intestinal peptide (VIP) and LAMP5/PAX6 (synonymous with HTR3A-expressing but lacking VIP) (*23*, *24*). Single cell transcriptomics studies of the mouse and human cortex revealed previously unrecognized diversity of neuron types within each of these four canonical interneuron subclasses (*1*, *25*, *26*). Mouse Patch-seq studies of cortical GABAergic neurons showed discrete morphological and electrophysiological properties between the subclasses yet a continuum within each subclass. (*16*, *17*) By comparison, similar studies of the human cortex have proven challenging due to the limited availability of adult human brain tissue and lack of genetic approaches to effectively target these spatially distributed and in some cases exceedingly rare cell types. Prior single neuron electrophysiology studies of human cortical GABAergic neurons are relatively few and have relied mainly on firing patterns (e.g., fast spiking (FS) vs. non-FS), limited marker genes, or gene panels to achieve approximate alignment to subclasses described in rodent (*3*, *27*–*32*). The previous lack of a comprehensive human cortical cell type taxonomy as a reference for these studies likely hindered efforts at classification, and the degree of conservation between human and mouse neocortical GABAergic cell type diversity and marker gene expression has only partially been delineated.

Here we combined rapid viral genetic labeling using an optimized DLX2.0 forebrain GABAergic enhancer viral vector (*33*) together with Patch-seq recordings in cortical slices derived from human neurosurgical resections. Mapping of recorded neurons to transcriptomically-defined types in our human cortical taxonomy facilitated a detailed functional annotation of human neocortical GABAergic cell subclasses and select abundant t-types. This work provides a promising roadmap for future functional studies of human brain cell types at the resolution of emerging transcriptomic cell type taxonomies and provides a rich open access dataset for exploring gene-function relationships in a wide diversity of human neocortical GABAergic neuron types.

## Results

We performed Patch-seq experiments to characterize the electrophysiological, morphological and transcriptomic profiles of GABAergic neurons in human neocortical brain slices using a well-established Patch-seq platform (*13*–*15*). To expand our sampling capability, both temporally, and to facilitate preferential targeting, we adopted an existing human *ex vivo* culture paradigm (*34*) to apply enhancer-based Adeno-Associated Virus (AAV) to cultured human neocortical slices (Fig. 1A) (*2*, *33*).

**Figure 1.**
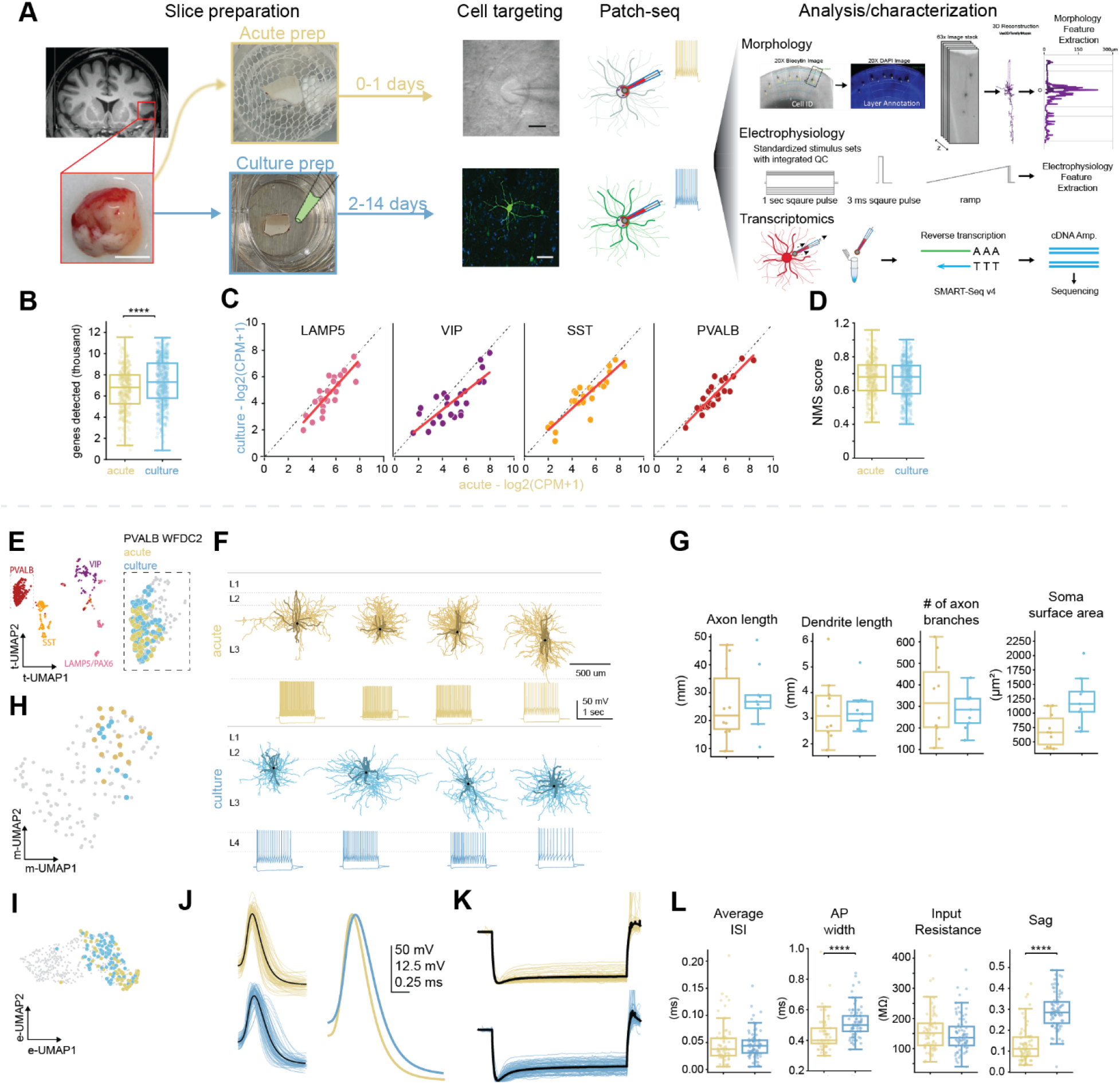
Slice culture paradigm and multimodal characterization. (A) Tissue processing schematic for acute and culture paradigm, patch-seq targeting guided by brightfield or fluorescence and subsequent multimodal analysis/characterization. Scale bars: human brain specimen – 500 um, brightfield of patched cell – 10 um, fluorescent image – 50 um. (B) Box plot represents the difference in the number of genes detected between the two paradigms. (C) Scatter plot of the average gene expression in acute versus culture for the top 25 differential expressed genes for the LAMP5/PAX6, VIP, SST and PVALB subclasses. Red line is regression line in each plot. (D) Box plot representing the difference in NMS score between the two paradigms. (E) UMAP representation of the PVALB subclass transcriptomic space with PVALB WFDC2 acute and culture cells highlighted. (F) Cortical depth matched PVALB WFDC2 morphologies from acute and culture shown aligned to an average cortical template, with corresponding voltage responses to a 1 s-long current step of −90 pA and rheobase +80 pA. (G) Box plots showing select morphology features for cortical depth matched PVALB WFDC2 from the acute and culture paradigm. (H) UMAP representation of morphology space with PVALB WFDC2 acute and culture cells highlighted. (I) UMAP representation of electrophysiology space with PVALB WFDC2 acute and culture cells highlighted. (J) Overlaid single action potential sweeps from acute and culture PVALB WFDC2 neurons, black lines represent the mean and overlaid to the right for direct comparison. (K) Overlaid and normalized voltage response to a −90 pA hyperpolarizing current step from acute and culture PVALB WFDC2 neurons. Black lines represent mean of the group (L) Box plots showing select, distinguishing, features from PVALB WFDC2 neurons from the acute and culture paradigm.

Human samples were obtained from neurosurgically resected neocortical tissues and processed with standardized protocols across three experimental sites in the U.S., Netherlands and Hungary (*14*, *15*, *18*, *35*); most samples originated from the temporal and frontal lobes, along with smaller fractions in parietal, occipital and cingulate areas (Data S1). Human Patch-seq neurons were mapped to a middle temporal gyrus (MTG) single nucleus transcriptomics-based reference taxonomy (*1*) and assigned to a transcriptomic subclass and type using a “tree mapping” classifier (see Methods) (*16*).

### Morphological, electrophysiological and transcriptomic features in the slice culture paradigm

The culture paradigm offers the unique ability to extend the viability of *ex vivo* human specimens/slices beyond the first day and the opportunity to apply enhancer-based AAVs for cell-type specific labeling. With the use of Patch-seq recordings we systematically examined the acute and culture brain slice paradigms with three different data modalities – transcriptome/gene expression, morphology, and electrophysiology. We found cells from Patch-seq experiments performed in culture preparations had a significantly higher number of genes detected compared to the acute preparations (Fig. 1B). This elevation was prevalent and remained consistent across all days sampled in culture (Fig. S1A). Recently it has been identified that some Patch-seq neurons may contain microglial markers (*36*) and we sought to determine if the increased number of genes we observed in the culture paradigm was specific to microglial markers. To do this we performed standard Seurat clustering on Patch-seq cells from the culture paradigm. These cells clustered into two domains, with or without microglial gene expression signatures, and each was independent of cell type (Fig. S1C, S1D). This microglial signature (Data S2) was also observed in a subset of cells collected using the acute paradigm, although the effect was significantly less pronounced (Fig. S1B and S1C). Importantly, despite the increased microglial marker detection, this did not preclude or hinder the ability to accurately map inhibitory subclasses (Fig. S1D). Upon examination of key differentially expressed genes responsible for inhibitory subclass discrimination, average gene expression was strongly correlated, demonstrating alignment between acute and culture preparations (Fig. 1C). Collectively, when examining the normalized marker sum (NMS) score, a method to quantify the expression of mapping-related marker genes (*14*, *37*), we found no significant difference between the acute and culture preparations (Fig. 1D). These results suggest that the key markers used to map cell types are not perturbed in the culture paradigm, sustaining the ability to accurately map cell types.

To further examine any differences between Patch-seq in acute and cultured slices, we performed an indepth investigation into a transcriptomically-defined type that was the most abundant type sampled – ‘Inhibitory L2-4 PVALB WFDC2’ (hereafter called PVALB WFDC2) across the two paradigms. The acute and culture datasets exhibited overlapping structure when visualizing the transcriptome via a Uniform Manifold Approximation Projection (UMAP) method (*38*) (Fig. 1E) demonstrating that the culture paradigm did not affect the ability to map cell types accurately. We also evaluated their morphological properties in culture (n= 9) and acute preparations (n=10) using cortical depth matched PVALB WFDC2 neurons (Fig. 1F). We assessed fifty of their morphological features and none differed significantly (Fig. S2A) (P > 0.05, false discovery rate (FDR)-corrected Mann–Whitney test). Additionally, dimensionality reduction of the 50 morphology features shows PVALB WFDC2 neurons from the culture and acute paradigms to occupy similar space in the PVALB rich portion of morphology UMAP, demonstrating consistency in the morphometric features between the two paradigms (Fig. 1H).

Analysis of electrophysiological responses to a standardized current clamp protocol revealed 43 of 84 features differed across the two paradigms (Fig. S2B) (P < 0.05, false discovery rate (FDR)-corrected Mann–Whitney test). Shown in Fig. 1J are overlaid voltage traces of a single action potential and response to hyperpolarizing current (Fig. 1K) with observable differences between the two paradigms from PVALB WFDC2. Action potential width and sag are two of the key features that were significantly different between the two groups (Fig. 1L); however, multiple other cardinal intrinsic and membrane properties, such as interspike interval or input resistance, were not. Despite the differences, when projecting all 84 electrophysiological features via the UMAP, both paradigms generally occupied a similar space (Fig. 1I), thus preserving the ability to classify type based on the electrophysiological features. Despite the shifts in select electrophysiological features, combined with consistency of morphological and transcriptomic features between acute versus culture paradigms, the ability to accurately characterize transcriptomic cell types was well preserved. Therefore, to generate a more comprehensive multimodal characterization of human interneurons, we have combined the data from acute and culture paradigms. This strategy provides the unique opportunity to conduct an in-depth examination of the phenotypic properties of transcriptomic subclasses and types of human GABAergic neurons.

### Comparison of features across human GABAergic subclasses

Using this paradigm, we were able to preferentially target GABAergic neurons in human *ex vivo* neocortical slices for Patch-seq experiments. We characterized the transcriptomic, intrinsic physiological, and/or morphological properties of 778 interneurons from the four canonical GABAergic subclasses in human cortex. We describe interneurons in cortical Layers 2-6 (L2-L6), across 44 transcriptomic types (Fig. S4). When visualized in Seurat-aligned transcriptomic space via UMAP, the Patch-seq dataset overlapped with snRNA-seq human MTG data set (*1*) demonstrating feasibility, consistency and accuracy of mapping quality (Fig. 2A). Visualizing electrophysiological (Fig. 2B) and morphological (Fig. 2C) features in UMAP space revealed PVALB neurons occupy one subregion of both electrophysiology and morphology UMAPs. While SST, VIP, and LAMP5/PAX6 are mostly found in the other subregion with distinct areas of enrichment for each. Proximity in the UMAP suggests quantitative multimodal features can be used to distinguish inhibitory subclasses.

**Figure 2.**
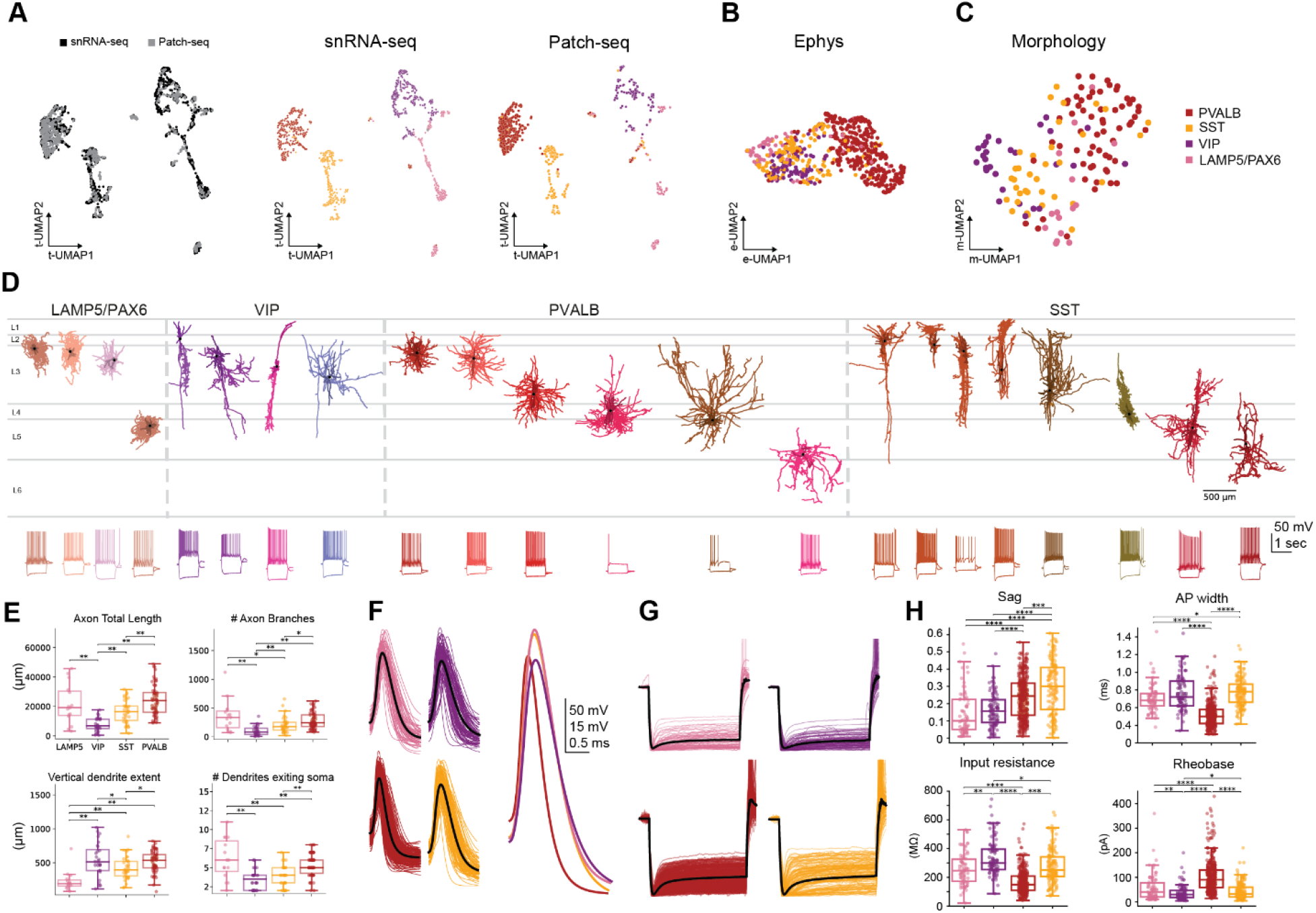
Human neocortical interneuron subclass characterization. (**A**) Integrated snRNAseq and patch-seq transcriptomic UMAP of inhibitory neurons. UMAP representations of electrophysiology (**B**) and morphology (**C**) space. (**D**) Representative morphologies from inhibitory subclasses with color distinguishing separate t-types shown aligned to an average cortical template with corresponding voltage responses to a 1 s-long current step to a −90 pA and rheobase +80 pA, shown below. (**E**) Box plots representing distinguishing morphology features by subclass. (**F**) Overlaid single action potential sweeps from each inhibitory subclass, black lines represents the mean and overlaid to the right for direct comparison. (**G**) Overlaid and normalized voltage response to a 1 sec −90 pA hyperpolarizing current step from each inhibitory subclass. Black lines represents mean of the group. (**H**) Box plots representing key, distinguishing electrophysiological features by subclass.

The dendrites and axons of a subset of neurons (n=140) with sufficient labeling was imaged at high resolution and reconstructed (Fig. 2D, Fig. S5). Neurons from the LAMP5/PAX6 subclass are dominated by classic neurogliaform morphologies. Although they are found in all cortical layers, we report predominately on neurons in L2 and L3. An in-depth investigation of L1 cells can be found in Chartrand et al. 2022. LAMP5/PAX6 neurons are distinguished by their stellate dendrites, numerous primary dendrites, and extensive axon branching. They also have the shortest soma to branch tip Euclidean distance for both axon and dendrite (Fig. 2E). The morphological dataset for the VIP subclass, also dominated by cells in L2 and L3, displays more diverse morphologies, many with bipolar dendrites and descending axon, either with a wide or narrow profile. The PVALB subclass, with multipolar dendrites, contains basket-like morphologies with radially arrayed axons that have a consistently wide horizontal span. A distinctive PVALB subclass neuron of type L4-5 MEPE (Fig. 2D, 4^th^ PVALB cell from the left) has axon that spans L3-4 and resembles mouse cortical interlaminar or translaminar basket cells (*16*, *39*).The SST subclass also contains diverse morphologies ranging from classical double bouquet cells to non-Martinotti cells. One notable human SST type described in Hodge et al. is L4-5 SST STK32A, which is the homolog of mouse Sst Hpse Cbln4 and Sst Hpse Sema3c (*1*). These homologous types share similar morphological qualities. Mouse Sst Hpse Cbln4 neurons are predominately found in L4 and have extensive axonal branching that strongly innervates L4 (*16*, *40*), which is similar to what we see for L4-5 SST STK32A neurons (Fig. 2D, 3^rd^ SST cell from the right). Interestingly, mouse Sst Hpse Cbln4 neurons are found to preferentially target L4 pyramidal cells (*41*), suggesting that they may also have selective connectivity in human.

Comprehensive analysis of the associated voltage traces for the four interneuron subclasses, LAMP5/PAX6 (n=75), VIP (n=99), SST (n=149), and PVALB (n=352), revealed distinct passive and active electrophysiological features (Fig. 2B, 2D, 2F-H), supporting previous studies (*23*, *42*). Example voltage traces are shown below the reconstructions in Fig. 2D and demonstrate diversity in intrinsic firing across the interneuron subclasses, ranging from adapting to irregular or fast-spinking. Shown in Fig. 2F are overlaid normalized voltage traces of a single action potential in response to a 3-ms current pulse with subclass average (right). Most notably, action potentials from the PVALB subclass have faster kinetics than the other subclasses, while LAMP5/PAX6 have a higher peak amplitude. Overlaid, normalized, voltage traces in response to hyperpolarizing current (Fig 2G) illustrate that LAMP5/PAX5 and VIP subclasses had minimal sag compared to PVALB and SST subclasses. Several additional features further demonstrate the electrophysiological distinctness of each subclass (Fig. 2H).

### Revising subclass assignment based on multimodal data

SST and PVALB subclasses are known to arise developmentally from the Medial Ganglionic Eminence (MGE) (*10*, *24*, *43*). Furthermore, the gene expression similarity between some SST and PVALB t-types additionally points to close links between these subclasses. For example, neurons mapping to ‘L2-4 SST FRZB’ (hereafter called SST FRZB), L2-4 PVALB WFDC2 or L4-6 PVALB SULF1 have intermediate relationships by expressing continuous variation in genes. (*1*). Using the previously described snRNA-seq dataset we identified the top 20 differentially expressed genes for the SST and PVALB subclasses and plotted expression across SST and PVALB t-types. From this analysis, we observed neurons belonging to the SST FRZB t-type notably lacked expression of multiple key genes that define each subclass (Fig. 3A). Given the intermediate nature of the SST FRZB gene expression pattern, we sought to gain a better understanding of the phenotypic properties of this t-type.

**Figure 3.**
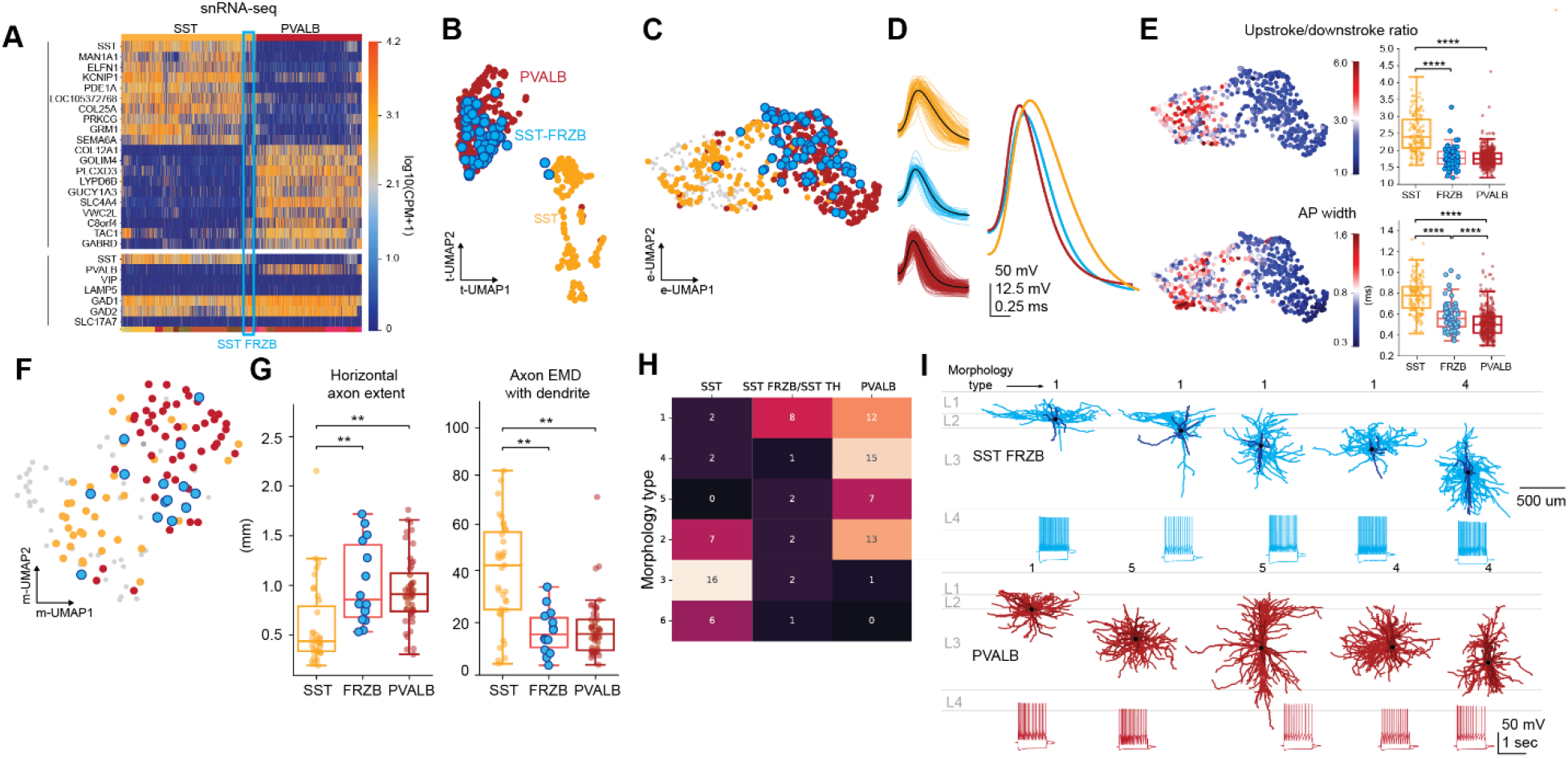
Phenotypic similarities of SST FRZB and the PVALB subclass. (**A**) Heat map of snRNA-seq data with the top 20 differentially expressed genes for SST and PVALB subclass with SST FRZB highlighted in blue. Genes in subpanel are key genes for the major subclasses and classes. (**B**) Transcriptomic UMAP highlighting the PVALB and SST subclasses and SST FRZB in blue. (**C**) UMAP representation of electrophysiology space with highlighting SST and PVLAB subclasses and SST FRZB in blue. (**D**) Overlaid single action potential sweeps from SST FRZB and PVALB subclass, black lines represent the mean and overlaid to the right for direct comparison. (**E**) UMAP representation of electrophysiology space color-coded by upstroke/downstroke ratio and action potential width. Box plots to the right show the distribution of data of the SST and PVALB subclass with SST FRZB highlighted in blue. **(F)** UMAP representation of morphology space with highlighting SST and PVLAB subclasses and SST FRZB in blue. **(G)** Morphology feature box plots for SST, SST FRZB and PVALB. **(H)** Morphology hierarchical co-cluster by SST, SST FRZB/SST TH and PVALB type. Numbers represent number of cells in each cluster. **(I)** Representative morphologies from SST FRZB and PVLAB subclass shown aligned to an average cortical template with associated voltage responses to a 1 s-long current step of −90 pA and rheobase +80 pA. Morphology classifications from 3H are shown above each reconstruction.

We took advantage of the multimodal approach of Patch-seq to examine how the phenotypic properties of SST FRZB are related to the PVALB and/or SST subclasses. First, we observed that the vast majority of SST FRZB samples localized to the PVALB subclass within the transcriptomic UMAP space in our Patch-seq dataset (Fig. 3B). Similarly, this was also observed with the snRNA-seq (data not shown). Phenotypically, SST FRZB was also most similar to the PVALB subclass. Examination of all 84 electrophysiological features, visualized using a UMAP, revealed that the majority of the SST FRZB neurons co-localized with the PVALB subclass (Fig. 3C). One defining feature of cortical PVALB interneurons is their fast-spiking behavior and narrow action potentials (*16*, *44*, *45*). We observed that the action potential kinetics, such as upstroke/downstroke ratio, action potential width and intrinsic firing of SST FRZB aligned with fast-spiking interneurons of the PVALB subclass (Fig. 3D, 3E, 3I).

Next, we investigated the morphological properties of SST FRZB neurons. We found that these neurons, when visualized in a morphological UMAP, overlap more with PVALB than with other SST neurons (Fig. 3F). Notably morphological features such as horizontal axon extent and a measure of the dissimilarity of the axon and dendrite compartments (axon/dendrite earth mover’s distance) also demonstrate that SST FRZB neurons have individual morphological features that are more similar to PVALB basket cells than SST neurons (Fig. 3G) (P >0.05, false discovery rate (FDR)-corrected). To test whether SST FRZB neurons would align with PVALB basket cells in an unbiased way we performed hierarchical co-clustering on all morphologies in the SST and PVALB subclasses. Clustering revealed three morphology types (1, 4, and 5) dominated by PVALB neurons and two types (3 and 6) dominated by neurons belonging to the SST subclass. We found that cells in SST FRZB and SST TH (a related, sparsely sampled deep layer SST t-type that also shows gene expression similarities to PVALB) more often clustered with PVALB morpho types (Fig. 3H). Juxtaposing morphologies of select SST FRZB neurons and PVALB cells also revealed their morphological similarity to each other, including multipolar dendrites and dense, local axon that avoids L1 (Fig. 3I). Based on these collective results we find that SST FRZB neurons should be considered as part of the PVALB subclass rather than SST subclass. This reclassification is applied for subclass level analyses throughout the manuscript and demonstrates the importance of collecting multimodal/functional data to validate transcriptomic cell types, particularly for cells with intermediate or ambiguous gene signatures.

### Morphological heterogeneity within SST CALB1

We identified dramatic morphological heterogeneity within the main supragranular SST t-type, SST CALB1. In mouse, SST subclass neurons in L2/3 have the classic Martinotti Cell (MC) shape, defined as having extensive L1 axon (*46*). In human, we see much more diversity. We qualitatively identified four morphological types within SST CALB1. Three of these types resembled previously described neurons: Martinotti Cells, Double Bouquet Cells (DBC), and Basket-like Cells (BC). We find classic Martinotti cells at the L2/3 border and in L3. The MCs contain considerable ascending axon resulting in an extensive plexus in L1 with some, comparatively minimal, descending axon. DBCs have a characteristic ratio of axon to dendrite width, which distinguishes these from other SST subclass neurons. We elaborate on DBCs, which are thought to be primate specialized, in a subsequent figure. We also find one SST CALB1 neuron with basket-like properties, including radially arrayed dendrite and axonal branches, and minimal L1 axon innervation. The fourth morphological type, a subset of the MCs, have unusually sparse axon compared to MC cells, and thus we refer to them as ‘sparse SST cells’ to distinguish them from classic MCs (Fig. 4A). Sparse SST neurons have somas restricted to L2, contain axons in L1 also with considerable descending axons with one instance of the axon reaching deep L5/upper L6. Unlike other morphology types within SST CALB1 this type has a striking wide dendrite and axonal extent, and sparsely branching axon that plateaus in L1 and L2 (see histograms Fig. 4A). Our dataset contains additional cells within the SST CALB1 t-type (Fig. S4) that we did not categorize into these four morphology types due to insufficient information to make a confident morphological qualification.

**Figure 4.**
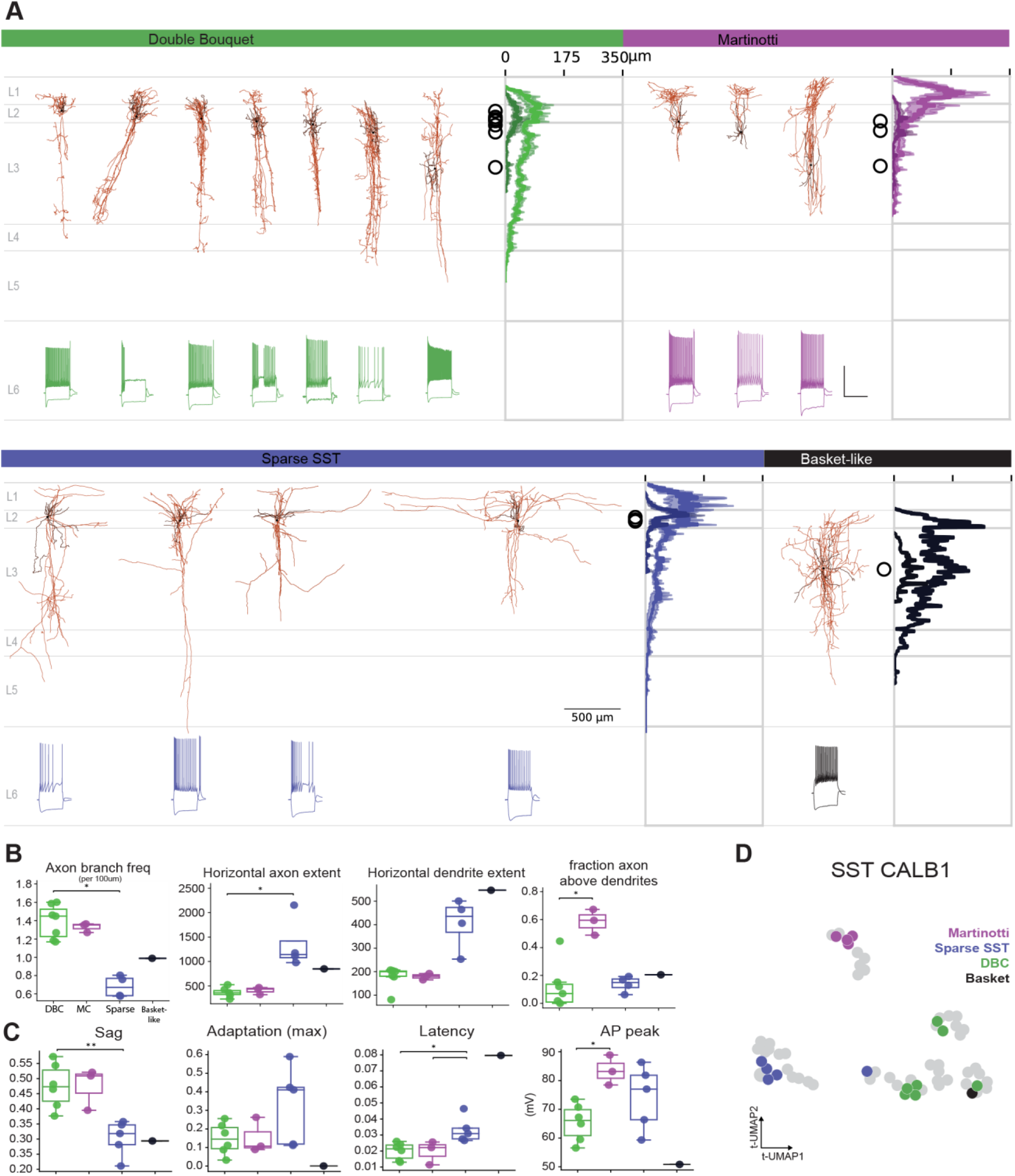
Morphological heterogeneity within SST CALB1. (**A**) Morphologies from SST CALB1 t-type categorized by qualitative morphology type shown aligned to an average cortical template, with histograms to the right of the morphologies displaying average dendrite (darker color) and axon (lighter color) branch length by cortical depth (shading shows +/− 1 SD about mean, soma locations represented by black circles). Voltage responses to a 1 s-long current step to a −90 pA and rheobase +80 pA, shown below. Box plots representing key morphological **(B)** and electrophysiological **(C)** features by morphology type. **(D)** UMAP of all SST CALB1 cells based on 253 genes differentially expressed between 10 DBC, MC and Sparse SST cells. Cells cluster into three main groups correspong to morphology types (colored markers). Cells with unknown morphologies are gray markers.

We next wanted to investigate how passive and active electrophysiological properties vary by the four morphological types identified in the SST CALB1 t-type. Despite the limited number of cells, we observed trends emerging that demonstrate how particular features may correspond with distinct morphologically defined SST CALB1 neurons. For example, Sparse SST cells show a lower degree of sag, a higher adaptation ratio and a slightly delayed on-set of action potential firing as compared to DBCs and MCs, while the peak of the action potential is highest in MCs (Fig. 4C). These data suggest a correspondence might exist between functional features and morphology for neurons within the heterogeneous SST CALB1 t-type.

We next sought to identify differentially expressed genes associated with the morphological heterogeneity within the SST CALB1 t-type and determine whether cells could independently be grouped into morphological types based on gene expression. Using this variable gene set, three distinct clusters appeared with morphological types largely separated across clusters (Fig. 4D). These results demonstrate that the morphological types we identified display distinct gene expression patterns and that even within a transcriptomic type, there are genetic correlates of differential phenotypic properties.

### Double bouquet cells

Most notably among the SST CALB1 morphological types is the double bouquet cell, specialized in primates (*47*, *48*) but also discovered in other specific carnivores (*49*–*51*). DBCs are described as having a “horse tail” morphology with ascending and descending narrow axon bundles. Though the morphology is well accepted in the field, the definition of molecular markers as well as their transcriptomic cell type identity remains elusive and incompletely resolved (*52*). Interestingly, we find these classical DBC morphologies map to two closely related t-types, SST CALB1, mentioned above, and SST ADGRG6. DBCs from both t-types have classic “horse tail” axon collaterals extending down to L5 and frequently up to L1. They often contain multiple descending axon bundles with short perpendicular branches. In addition to a narrow axon signature, we find their dendrites to be narrow, and either bi-tufted or multipolar (Fig. 5A). MERFISH analysis to map the spatial distributions of all types in the MTG taxonomy reveals the SST CALB1 t-type is fairly restricted to L2 while SST ADGRG6 is more broadly distributed across L2 and 3 (Fig. 5B). In agreement with the differing soma distribution patterns, we see a slight shift in their axon peak distribution with SST CALB1 axon shifted more superficially than SST ADGR6 (histograms at right, Fig. 5A). We also found that 6 out of 7 DBCs in the SST CALB1 t-type were derived from temporal cortex specimens whereas 3 out of 4 DBCs in the SST ADGRG6 t-type were from frontal cortex specimens, suggesting a possible differential abundance or enrichment across the cortex (Fig. S3A). Therefore, we observed two distinct populations of human DBCs that are phenotypically virtually indistinguishable at present but that show different areal enrichment and map to distinct SST t-types occupying distinct laminar positions in the cortical depth.

**Figure 5.**
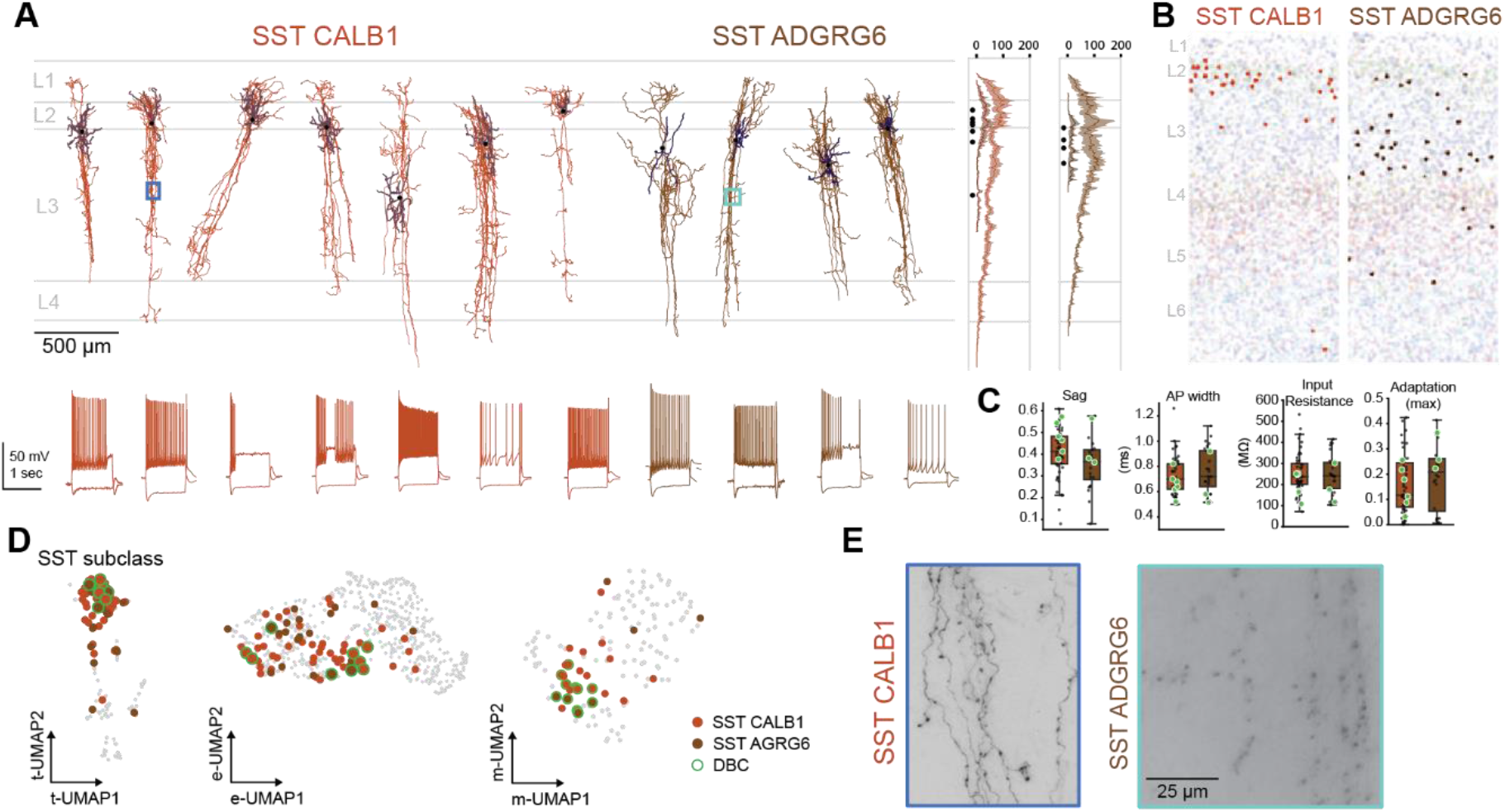
Multimodal properties of Double Bouqeut Cells. **(A)** Putative DBCs morphologies map to transcriptomic type SST CALB1 or SST ADGRG6, shown aligned to an average cortical template. Histograms to the right of the morphologies displaying average dendrite (darker color) and axon (lighter color) branch length by cortical depth (shading shows +/− 1 SD about mean, soma locations represented by black circles). Accompaning each reconstruction is the voltage response to a 1 s-long current step to a −90 pA and rheobase +80 pA, for each respective DBC cell (bottom). **(B)** Spatial distribution fo SST CALB1 and SST ADGRG6 revealed by MERFISH. **(C)** 4 specific electrophysiology features shown as box/scatter plots for the SST CALB1 and SST ADGRG6 t-types with putative DBC highlighted in green.**(D)** UMAP representation of transcriptpomics (isolated to the SST subclass), electrohpysiology and morphology. SST CALB1 (orange) and SST ADGRG6 (brown) t-types and qualitatively defined putative Double Bouquet Cells (DBC) highlighted in green. **(E)** 63x MIP inset corresponding to 4A showing axon “horsetail” bundles and boutons.

Given that we find DBCs within two t-types and taking into account a recent report of variable or inconsistent gene expression signatures for morphologically-defined human DBCs (*52*), we next sought to explore how these results compared to marker gene expression for human DBCs in our dataset. As shown in Fig. S3A, most DBCs show strong expression of CALB1 (Calbindin) while only one shows expression of CALB2 (Calretinin). All but one cell shows some degree of SST expression and mixed expression of TAC1, NOS1 and NPY (Fig. S3A). With this variable expression and limited sample size, there are no key genes specific to morphologically-defined DBCs.

To examine the functional properties of DBCs we examined the voltage responses to hyperpolarizing and depolarizing current injection. DBCs display a range of firing patterns from accommodating and stuttering to irregular spiking (Fig. 5C). Furthermore, in the electrophysiological UMAP space, DBCs from SST CALB1 and SST ADGRG6 types are intermingled with other SST cells (Fig. 5D, middle). Representative electrophysiological features show broad distribution of individual DBCs across the two t-types. One main feature that was consistent and prominent with all DBCs was a higher sag ratio (Fig. 4C and 5E), in close agreement with the recently reported electrophysiological signature of human DBCs (*52*).

In morphological-UMAP space, we see DBCs from these two t-types cluster near each other with perhaps some separation at the t-type level (Fig. 5D, right). Bouton density measurements suggest SST ADRG6 is slightly denser than SST CALB1 although this does not reach significance (Fig. 5E and S3C). The DBCs from these two t-types largely overlap in their morpho-electric properties, and their Patch-seq soma distributions align with MERFISH, suggesting that this morphological type with different depth distributions belong to the different t-types, SST CALB1 and SST ADGR6.

## Discussion

The remarkable diversity of cortical interneurons has posed a major challenge to neuroscientists seeking to classify and characterize their defining properties since the dawn of cellular neuroscience. In recent years, the emergence of single cell transcriptomics with high depth of sampling of cells or nuclei has served as a powerful and scalable approach to derive the complete parts list of a given brain region or diverse bodily organs and tissues. Our work leveraged the Allen Institute human MTG cell type taxonomy (*1*) as a reference to map the transcriptomic identities of patch-clamp recorded human cortical GABAergic neurons in *ex vivo* brain slices. This taxonomy contains 45 GABAergic neuron t-types across four canonical interneuron subclasses (LAMP5/PAX6, VIP, SST, and PVALB). In the present study, we provide an important conceptual advance of combining viral genetic labeling with Patch-seq mapping to prospectively label and target human cortical GABAergic neurons versus the far more abundant glutamatergic neuron types. We performed Patch-seq experiments on 778 human cortical GABAergic neurons and aggregated our data at the subclass level to uncover the signature morphoelectric properties of these canonical human interneuron subclasses, and where feasible, a deeper dive into features of select abundant GABAergic t-types. Although many of the GABAergic t-types were not sampled at sufficient depth to conclude their defining properties in this study, remarkably our dataset achieved coverage of 44 of the 45 GABAergic neuron t-types as found in the reference taxonomy, many of which represent rare cell types. Thus, with adequate time and sustained effort, a comprehensive Patch-seq sampling and quantitative analysis of all human cortical GABAergic neurons is now within reach.

Importantly, we characterized the changes that occur in this short-term slice culture and viral labeling paradigm in multiple data modalities including gene expression, electrophysiological and morphological features (where sampling depth permitted). Although discrete changes were evident and described in electrophysiology (e.g., most notably increased sag), these putative culture differences did not preclude or hinder high confidence mapping of recorded neurons to GABAergic subclasses and t-types in the human MTG taxonomy. This is well explained by the fact that marker genes important for mapping at the GABAergic subclass level were robustly detected overall and minimally impacted by slice culture or viral transduction. In contrast, we identified a notable gene expression signature in a subset of our dataset corresponding to a module of microglia related genes. Surprisingly, this identical signature was observed for both neurons recorded in acute and cultured slices (but more pronounced in culture) and has been similarly observed in other human and mouse acute brain slice Patch-seq datasets (*15*–*17*, *35*, *36*). Further work is warranted to understand the cause and consequences of this gene expression signature in brain slice experiments. Notwithstanding, the short-term brain slice culture and viral genetic labeling paradigm proved instrumental for extending the window of opportunity and improving efficiency of sampling GABAergic neuron types in precious human brain tissue samples.

A comparison of viral genetic targeting versus unbiased Patch-seq experiments revealed a significant increase in coverage of neurons in the SST subclass and allowed us to aggregate a sizable dataset of neurons mapping to the SST CALB1 t-type. These neurons exhibited unexpected heterogeneity in morphoelectric features that is seemingly incongruent with the mapping outcome to a single t-type in the taxonomy. At least three discrete morphological types were recognized including DBCs with classical descending long “horse tail” axon bundles (*53*), Martinotti cells with ascending axons reaching L1 (*46*), and a type we labeled sparse SST for the sparsely branched and lengthy dendrite segments. Further grouping by morpho type within this SST CALB1 t-type revealed strong coherence of electrophysiological features with many distinctive features between groups. These results suggest that further splitting of the SST CALB1 t-type may be warranted but will require additional studies to fully resolve. On a similar note, although we were underpowered in our differential gene expression analysis between these morpho defined groupings, we make these gene lists available to stimulate new experiments or analysis that could help uncover salient functional differences.

Another important outcome from our study is the delineation of the transcriptomic cell types containing human DBCs. In addition to SST CALB1 t-type mapping, we also observed DBCs mapping to a second t-type in the SST subclass, SST ADGRG6. Interestingly, DBCs mapping to the SST CALB1 t-type were predominantly found in temporal cortex samples, whereas DBCs mapping to the SST ADGRG6 t-type were enriched in frontal cortex samples, suggesting possible regional differences in abundance of these putative DBC subtypes across the human cortex. Furthermore, our spatial transcriptomics analysis revealed a clear shift in laminar distribution between the two t-types, with SST CALB1 soma distribution restricted to L2 and upper L3 and SST ADGRG6 soma distribution extending into L5. DBCs sampled from Patch-seq experiments follow this trend of distinct laminar positioning of the two t-types. We were not able to discriminate DBCs vs non-DBCs within these t-types using current available spatial methodologies. Nonetheless, these data are intriguing and consistent with the existence of at least two subtypes of human DBCs. Whether these transcriptomically defined subtypes align to previously reported DBC subtypes in human cortex (*51*), or if there exist functional specializations between these subtypes remains to be determined. Our findings on human DBCs corroborate and extend recent work by Somogyi and colleagues (*52*) that detailed the electrophysiological signatures of this enigmatic human cell type and provided important clues about their molecular identity, synaptic connectivity, and function. The overall picture from these two independent studies is that human DBCs may be more heterogeneous than previously suspected in terms of molecular markers. Mapping DBCs based on single cell transcriptomes versus analysis of small panels of antibodies or RNA probes allowed us to overcome confounds related to false negative detection (e.g., lack of SST expression detected in the prior study) and to definitively annotate SST subclass membership and pin down the two t-types containing this morphological cell type in the human MTG cell type taxonomy.

Our dataset has a notable limited sampling of neurons with classic Martinotti morphologies in the SST subclass. This may be the result of biased sampling of supragranular layers in our experiments, whereas SST+ Martinotti cells could instead be more abundant in infragranular layers of the human neocortex. The abundance and spatial distribution of Martinotti cells in the human cortex has not been detailed to our knowledge. The upper layer sampling bias was evident in both unlabeled and virus labeled slices. We observed denser GABAergic neuron labeling in L2 and L3 with our optimized DLX2.0 enhancer virus, suggesting possible enrichment for GABAergic neuron types most abundant in these layers (e.g., SST CALB1). However, other neuron types with expected enrichment in supragranular layers such as chandelier cells of the PVALB subclass and many diverse t-types of the VIP subclass were poorly sampled in this study. The underlying explanation remains unclear, but one possibility may be unexpected laminar shifts in these cell types in human versus rodent cortex. For example, chandelier cells are known to be enriched at the L1/2 border in rodent cortex (*54*) but instead show enrichment at the L3/4 border in human temporal cortex (*1*). Rapid progress in single cell epigenetic data generation (*55*–*58*) and continued discovery of novel brain cell type specific enhancers suitable for use in viral vectors (*33*, *59*–*61*) could be the key to targeted analysis of these and other important human cortical interneuron types in the future.

Perhaps the most impactful finding of our study is the direct demonstration of how multimodal Patch-seq data is vital to refinement of transcriptomic cell type taxonomies. Our findings clearly support a reclassification of the SST FRZB t-type from the SST subclass to the PVALB subclass in our human MTG taxonomy. This confusion was no doubt the result of ambiguous expression (or lack thereof) of numerous SST and PVALB subclass marker genes in the SST FRZB t-type and perhaps also rooted in the shared medial ganglionic eminence developmental origin of SST and PVALB subclasses (*43*). This reclassification underscores the notion that cellular taxonomies built on single cell transcriptomes and differential gene expression are not static, but rather represent a starting foundation to build upon as new data modalities are obtained and aligned at the resolution of t-types. Thus, transcriptomic cell type taxonomies are necessary but not always sufficient to infer meaningful functional types with high accuracy. Alignment of multimodal data such as spatial distribution and abundance, cellular morphology, axonal projections and connectivity, neuromodulation, intrinsic and synaptic electrophysiological properties will be essential to refine and extend foundational cellular taxonomies of the brain. Examples of refinement could include reclassification of types (e.g., SST FRZB t-type), splitting of types (e.g., heterogeneous properties of SST CALB1 t-type), or merging of types with graded or high similarity of features or functions in brain circuits (yet to be determined). In this study we have provided a first order characterization of the signature morphoelectric properties of the canonical human cortical GABAergic neuron subclasses and select t-types and a rich open-access Patch-seq dataset for further exploration of gene-function relationships and comparative studies of homologous cell types across species.

## Author Contributions

Conceptualization – BRL, RD, BK, SAS, EL, JTT

Data Curation - BRL, RD, JAM, TC, JC, RM, AM, LN, EJM, JN, LP, CDK, KS, JW

Formal analysis - BRL, RD, JAM, TC, RM, AM, LN, NJ, NJ, NW, TB, NWG

Funding acquisition - BL, NAG, HDM, GT, EL, JTT

Investigation - BRL, RD, JAM, TC, JC, RM, AM, LN, LA, KB, DB, KB, TC, EAC, ND, ND, SLWD, TE, RE, AAG, AG, EG, JG, KH, TSH, DH, LK, AKK, MK, BL, DM, GM, XO, AP, LP, RR, AR, IS, MT, JT, SV, FW, NW, GW, JW, RH, JTT

Methodology - BRL, RD, JAM, JC, RM, AM, LN, DB, KB, TC, ND, ND, MK, BL, MM, TB, RH, BL, BK, SAS, JTT

Project administration - BRL, RD, EJM, JN, SMS, BT, DV, LE, KS, JW, JB, NAG, BK, CPJdK, LK, HDM, SAS, GT, EL, JTT

Resources - BRL, RD, ND, EJM, MTM, DN, XO, NT, NW, SY, PB, CC, RGE, MF, NWG, BG, RPG, JSH, CDK, ALK, JGO, AP, JR, DLS, KS, JW, EL, JTT

Software - TC, MM, NWG, TJ

Supervision - BRL, RD, JC, ND, MM, DM, LP, SY, TB, TJ, KS, JW, HZ, JB, NAG, BK, CPJdK, HDM, SAS, GT, EL, JTT

Visualization - BRL, RD, JAM, TC, JC, RM, AM, LN, NJ, NJ, MM, NWG

Writing - Original Draft – BRL, RD, JAM, JTT

Writing – Review & Editing - BRL, RD, JAM, TC, BK, CPJdK, HDM, SAS, GT, EL, JTT

## Data and materials availability

All data from the study will be made available through BIL, DANDI, NeMO repositories. Other software libraries essential to the analysis are publicly available at https://github.com/AllenInstitute/patchseqtools (Patch-seq), https://github.com/alleninstitute/ipfx (electrophysiology), https://github.com/AllenInstitute/neuron_morphology and https://github.com/AllenInstitute/skeleton_keys (morphology), and https://github.com/AllenInstitute/scrattch/ (transcriptomics).

## Materials and Methods

### Human tissue acquisition

Surgical specimens were obtained from local hospitals (Harborview Medical Center, Swedish Medical Center and University of Washington Medical Center) in collaboration with local neurosurgeons. Data included in this study were obtained from neurosurgical tissue resections for the treatment of refractory temporal lobe epilepsy or deep brain tumor. All patients provided informed consent and experimental procedures were approved by hospital institute review boards before commencing the study. Tissue was placed in slicing artificial cerebral spinal fluid (ACSF) as soon as possible following resection. Slicing ACSF comprised (in mM): 92 *N*-methyl-D-glucamine chloride (NMDG-Cl), 2.5 KCl, 1.2 NaH_2_PO_4_, 30 NaHCO_3_, 20 4-(2-hydroxyethyl)-1-piperazineethanesulfonic acid (HEPES), 25 D-glucose, 2 thiourea, 5 sodium-L-ascorbate, 3 sodium pyruvate, 0.5 CaCl_2_.4H_2_O and 10 MgSO_4_.7H_2_O. Before use, the solution was equilibrated with 95% O_2_, 5% CO_2_ and the pH was adjusted to 7.3 by addition of 5N HCl solution. Osmolality was verified to be between 295–305 mOsm kg^-1^. Human surgical tissue specimens were immediately transported (15–35 min) from the hospital site to the laboratory for further processing.

### Human neurosurgical specimens and ethical compliance

The neurosurgical tissue specimens collected for this study were apparently non-pathological tissues removed during the normal course of surgery to access underlying pathological tissues. Tissue specimens were determined to be non-essential for diagnostic purposes by medical staff and would have otherwise been discarded. Tissue procurement from neurosurgical donors was performed outside of the supervision of the Allen Institute at a local hospital and tissue was provided to the Allen Institute under the authority of the institutional review board of the participating hospital. A hospital-appointed case coordinator obtained informed consent from the donor before surgery. Tissue specimens were deidentified before receipt by Allen Institute personnel.

### Tissue processing

Human acute and cultured brain slices (350 μm) were prepared with a Compresstome VF-300 (Precisionary Instruments) modified for block-face image acquisition (Mako G125B PoE camera with custom integrated software) before each section to aid in registration to the common reference atlas. Brains or tissue blocks were mounted to preserve intact pyramidal neuron apical dendrites within the brain slice. Slices were transferred to a carbogenated (95% O_2_/5% CO_2_) and warmed (34 °C) slicing ACSF to recover for 10 min according to the NMDG protective recovery method (*62*). Acute brain slices were then transferred to room temperature holding ACSF of the composition (in mM): 92 NaCl, 2.5 KCl, 1.2 NaH_2_PO_4_, 30 NaHCO_3_, 20 HEPES, 25 D-glucose, 2 thiourea, 5 sodium-L-ascorbate, 3 sodium pyruvate, 2 CaCl_2_.4H_2_O and 2 MgSO_4_.7H_2_O for the remainder of the day until transferred for patch clamp recordings. Before use, the solution was equilibrated with 95% O_2_, 5% CO_2_ and the pH was adjusted to 7.3 using NaOH. Osmolality was verified to be between 295–305 mOsm kg^-1^. Alternately, slices for interface culture were placed onto membrane inserts (Millipore) in 6 well plates with 1 mL per well of slice culture media of the composition: 8.4 g/L MEM Eagle medium, 20% heat-inactivated horse serum, 30 mM HEPES, 13 mM d-glucose, 15 mM NaHCO3, 1 mM ascorbic acid, 2 mM MgSO4·7H2O, 1 mM CaCl2.4H2O, 0.5 mM GlutaMAX-I and 1 mg/L insulin. The slice culture medium was carefully adjusted to pH 7.2–7.3 and osmolality of 300–310 mOsmoles per kilogram by addition of pure H2O, sterile-filtered and stored at 4°C for up to 2 weeks. Culture plates were placed in a humidified 5% CO2 incubator at 35°C, and the slice culture medium was replaced every two to three days until endpoint analysis. One to three hours after brain slices were plated on cell culture inserts, brain slices were infected by direct application of concentrated AAV viral particles over the slice surface (*34*).

### Patch clamp recording

Slices were continuously perfused (2 mL/min) with fresh, warm (34 °C) recording ACSF containing the following (in mM): 126 NaCl, 2.5 KCl, 1.25 NaH_2_PO_4_, 26 NaHCO_3_, 12.5 D-glucose, 2 CaCl_2_.4H2O and 2 MgSO_4_.7H2O (pH 7.3) and continuously bubbled with 95 % O2/5 % CO2. The bath solution contained blockers of fast glutamatergic (1 mM kynurenic acid) and GABAergic synaptic transmission (0.1 mM picrotoxin). Thick-walled borosilicate glass (Warner Instruments, G150F-3) electrodes were manufactured (Narishige PC-10) with a resistance of 4–5 MΩ. Before recording, the electrodes were filled with ~1.0–1.5 μL of internal solution with biocytin (110 mM potassium gluconate, 10.0 mM HEPES, 0.2 mM ethylene glycol-bis (2-aminoethylether)-N,N,N’,N’tetraacetic acid, 4 mM potassium chloride, 0.3 mM guanosine 5’-triphosphate sodium salt hydrate, 10 mM phosphocreatine disodium salt hydrate, 1 mM adenosine 5’-triphosphate magnesium salt, 20 μg/mL glycogen, 0.5 U/μL RNAse inhibitor (Takara, 2313A) and 0.5 % biocytin (Sigma B4261), pH 7.3). The pipette was mounted on a Multiclamp 700B amplifier headstage (Molecular Devices) fixed to a micromanipulator (PatchStar, Scientifica).

The composition of bath and internal solution as well as preparation methods were chosen to maximize the tissue quality of slices from adult mice, to align with solution compositions typically used in the field (to maximize the chance of comparison to previous studies), modified to reduce RNAse activity and ensure maximal gain of mRNA content.

Electrophysiology signals were recorded using an ITC-18 Data Acquisition Interface (HEKA). Commands were generated, signals processed, and amplifier metadata were acquired using MIES written in Igor Pro (Wavemetrics). Data were filtered (Bessel) at 10 kHz and digitized at 50 kHz. Data were reported uncorrected for the measured (Neher, 1992)–14 mV liquid junction potential between the electrode and bath solutions.

Prior to data collection, all surfaces, equipment, and materials were thoroughly cleaned in the following manner: a wipe down with DNA away (Thermo Scientific), RNAse Zap (Sigma-Aldrich), and finally nuclease-free water.

After formation of a stable seal and break-in, the resting membrane potential of the neuron was recorded (typically within the first minute). A bias current was injected, either manually or automatically using algorithms within the MIES data acquisition package, for the remainder of the experiment to maintain that initial resting membrane potential. Bias currents remained stable for a minimum of 1 s before each stimulus current injection.

To be included in analysis, a neuron needed to have a > 1 GΩ seal recorded before break-in and an initial access resistance <20 MΩ and <15 % of the Rinput. To stay below this access resistance cut-off, neurons with a low input resistance were successfully targeted with larger electrodes. For an individual sweep to be included, the following criteria were applied: (1) the bridge balance was <20 MΩ and <15 % of the Rinput; (2) bias (leak) current 0 ± 100 pA; and (3) root mean square noise measurements in a short window (1.5 ms, to gauge high frequency noise) and longer window (500 ms, to measure patch instability) were <0.07 mV and 0.5 mV, respectively.

Upon completion of electrophysiological examination, the pipette was centered on the soma or placed near the nucleus (if visible). A small amount of negative pressure was applied (~−30 mbar) to begin cytosol extraction and attract the nucleus to the tip of the pipette. After approximately one minute, the soma had visibly shrunk and/or the nucleus was near the tip of the pipette. While maintaining the negative pressure, the pipette was slowly retracted in the x and z direction. Slow, continuous movement was maintained while monitoring pipette seal. Once the pipette seal reached >1 GΩ and the nucleus was visible on the tip of the pipette, the speed was increased to remove the pipette from the slice. The pipette containing internal solution, cytosol, and nucleus was removed from the pipette holder and contents were expelled into a PCR tube containing lysis buffer (Takara, 634894).

### Transcriptomic Data Collection

#### cDNA amplification and library construction

We used the SMART-Seq v4 Ultra Low Input RNA Kit for Sequencing (Takara, 634894) to reverse transcribe poly(A) RNA and amplify full-length cDNA according to the manufacturer’s instructions. We performed reverse transcription and cDNA amplification for 20 PCR cycles in 0.65 ml tubes, in sets of 88 tubes at a time. At least one control eight-strip was used per amplification set, which contained four wells without cells and four wells with 10 pg control RNA. Control RNA was either Universal Human RNA (UHR) (Takara 636538) or control RNA provided in the SMART-Seq v4 kit. All samples proceeded through Nextera XTDNA Library Preparation (Illumina FC-131–1096) using either Nextera XT Index Kit V2 Sets A-D (FC-131–2001, 2002, 2003, 2004) or custom dual-indexes provided by IDT (Integrated DNA Technologies). Nextera XT DNA Library prep was performed according to manufacturer’s instructions, except that the volumes of all reagents including cDNA input were decreased to 0.2 x by volume. Each sample was sequenced to approximately 500 k reads.

#### RNA-sequencing

Fifty-base-pair paired-end reads were aligned to GRCm38 (mm10) using a RefSeq annotation gff file retrieved from NCBI on 18 January 2016 (https://www.ncbi.nlm.nih.gov/genome/annotation_euk/all). Sequence alignment was performed using STAR v2.5.3 (*63*) in two pass Mode. PCR duplicates were masked and removed using STAR option ‘bamRemoveDuplicates’. Only uniquely aligned reads were used for gene quantification. Gene counts were computed using the R Genomic Alignments package (*64*). Overlaps function using ‘IntersectionNotEmpty’ mode for exonic and intronic regions separately. Exonic and intronic reads were added together to calculate total gene counts; this was done for both the reference dissociated cell data set and the Patch-seq data set of this study.

#### SMART-seq v4 RNA-sequencing

The SMART-Seq v4 Ultra Low Input RNA Kit for Sequencing (Takara #634894) was used per the manufacturer’s instructions. Standard controls were processed with each batch of experimental samples as previously described. After reverse transcription, cDNA was amplified with 21 PCR cycles. The NexteraXT DNA Library Preparation (Illumina FC-131-1096) kit with NexteraXT Index Kit V2 Sets A-D (FC-131-2001, 2002, 2003, or 2004) was used for sequencing library preparation. Libraries were sequenced on an Illumina HiSeq 2500 instrument (Illumina HiSeq 2500 System, RRID:SCR_016383) using Illumina High Output V4 chemistry. The following instrumentation software was used during data generation workflow; SoftMax Pro v6.5; VWorks v11.3.0.1195 and v13.1.0.1366; Hamilton Run Time Control v4.4.0.7740; Fragment Analyzer v1.2.0.11; Mantis Control Software v3.9.7.19.

#### SMART-seq v4 gene expression quantification

Raw read (fastq) files were aligned to the GRCh38 human genome sequence (Genome Reference Consortium, 2011) with the RefSeq transcriptome version GRCh38.p2 (RefSeq, RRID:SCR_003496, current as of 4/13/2015) and updated by removing duplicate Entrez gene entries from the gtf reference file for STAR processing. For alignment, Illumina sequencing adapters were clipped from the reads using the fastqMCF program (from ea-utils). After clipping, the paired-end reads were mapped using Spliced Transcripts Alignment to a Reference (STAR v2.7.3a, RRID:SCR_015899) using default settings. Reads that did not map to the genome were then aligned to synthetic construct (i.e. ERCC) sequences and the E. coli genome (version ASM584v2). Quantification was performed using summerizeOverlaps from the R package GenomicAlignments v1.18.0. Expression levels were calculated as counts per million (CPM) of exonic plus intronic reads.

### Anatomical Annotations

#### Layer annotation and alignment

To characterize the position of biocytin-labeled cells, a 20× brightfield and fluorescent image of DAPI (4’,6-diamidino-2-phenylindole) stained tissue was captured and analyzed to determine layer position. Using the brightfield and DAPI image, soma position and laminar borders were manually drawn for all neurons and were used to calculate depth relative to the pia, white matter, and/or laminar boundaries. Laminar locations were calculated by finding the path connecting pia and white matter that passed through the cell’s soma coordinate, and measuring distance along this path to laminar boundaries, pia and white matter.

For reconstructed neurons, laminar depths were calculated for all segments of the morphology, and these depths were used to create a “layer-aligned” morphology by first rotating the pia-to-WM axis to vertical, then projecting the normalized laminar depth of each segment onto an average cortical layer template.

#### Human brain region pinning

Available surgical photodocumentation (MRI or brain model annotation) is used to place the human tissue blocks in approximate 3D space by matching the photodocumentation to a MRI reference brain volume “ICBM 2009b Nonlinear Symmetric”(*65*), with Human CCF overlayed (*66*) within the ITK-SNAP interactive software.

### Morphological Reconstruction

#### Biocytin histology

A horseradish peroxidase (HRP) enzyme reaction using diaminobenzidine (DAB) as the chromogen was used to visualize the filled cells after electrophysiological recording, and 4,6-diamidino-2-phenylindole (DAPI) stain was used to identify cortical layers as described previously (*15*).

#### Imaging of biocytin-labelled neurons

Mounted sections were imaged as described previously (*45*). In brief, operators captured images on an upright AxioImager Z2 microscope (Zeiss, Germany) equipped with an Axiocam 506 monochrome camera and 0.63× Optivar lens. Two-dimensional tiled overview images were captured with a 20× objective lens (Zeiss Plan-NEOFLUAR 20×/0.5) in bright-field transmission and fluorescence channels. Tiled image stacks of individual cells were acquired at higher resolution in the transmission channel only for the purpose of automated and manual reconstruction. Light was transmitted using an oil-immersion condenser (1.4 NA). High-resolution stacks were captured with a 63× objective lens (Zeiss Plan-Apochromat 63×/1.4 Oil or Zeiss LD LCI Plan-Apochromat 63x/1.2 Imm Corr) at an interval of 0.28 μm (1.4 NA objective) or 0.44 μm (1.2 NA objective) along the z axis. Tiled images were stitched in ZEN software and exported as single-plane TIFF files.

#### Morphological reconstruction

Reconstructions of the dendrites and the full axon were generated for a subset of neurons with good quality transcriptomics, electrophysiology and biocytin fill. Reconstructions were generated based on a 3D image stack that was run through a Vaa3D-based image processing and reconstruction pipeline (Peng et al., 2010). For some cells images were used to generate an automated reconstruction of the neuron using TReMAP (Zhou et al., 2016). Alternatively, initial reconstructions were created manually using the reconstruction software PyKNOSSOS (https://www.ariadne.ai/) or the citizen neuroscience game Mozak (Roskams and Popović, 2016) (https://www.mozak.science/). Automated or manually-initiated reconstructions were then extensively manually corrected and curated using a range of tools (for example, virtual finger and polyline) in the Mozak extension (Zoran Popovic, Center for Game Science, University of Washington) of Terafly tools (*67*, *68*) in Vaa3D. Every attempt was made to generate a completely connected neuronal structure while remaining faithful to image data. If axonal processes could not be traced back to the main structure of the neuron, they were left unconnected.

### MERFISH data generation

Human postmortem frozen brain tissue was embedded in Optimum Cutting Temperature medium (VWR,25608-930) and sectioned on a Leica cryostat at −17 C at 10 um onto Vizgen MERSCOPE coverslips. These sections were then processed for MERSCOPE imaging according to the manufacturer’s instructions. Briefly: sections were allowed to adhere to these coverslips at room temperature for 10 min prior to a 1 min wash in nuclease-free phosphate buffered saline (PBS) and fixation for 15 min in 4% paraformaldehyde in PBS. Fixation was followed by 3×5 minute washes in PBS prior to a 1 min wash in 70% ethanol. Fixed sections were then stored in 70% ethanol at 4C prior to use and for up to one month. Human sections were photobleached using a 150W LED array for 72 h at 4C prior to hybridization then washed in 5 ml Sample Prep Wash Buffer (VIZGEN 20300001) in a 5 cm petri dish. Sections were then incubated in 5 ml Formamide Wash Buffer (VIZGEN 20300002) at 37C for 30 min. Sections were hybridized by placing 50ul of VIZGEN-supplied Gene Panel Mix onto the section, covering with parafilm and incubating at 37 C for 36-48 h in a humidified hybridization oven. Following hybridization, sections were washed twice in 5 ml Formamide Wash Buffer for 30 minutes at 47 C. Sections were then embedded in acrylamide by polymerizing VIZGEN Embedding Premix (VIZGEN 20300004) according to the manufacturer’s instructions. Sections were embedded by inverting sections onto 110 ul of Embedding Premix and 10% Ammonium Persulfate (Sigma A3678) and TEMED (BioRad 161-0800) solution applied to a Gel Slick (Lonza 50640) treated 2×3 glass slide. The coverslips were pressed gently onto the acrylamide solution and allowed to polymerize for 1.5h. Following embedding, sections were cleared for 24-48 h with a mixture of VIZGEN Clearing Solution (VIZGEN 20300003) and Proteinase K (New England Biolabs P8107S) according to the Manufacturer’s instructions. Following clearing, sections were washed twice for 5 min in Sample Prep Wash Buffer (PN 20300001). VIZGEN DAPI and PolyT Stain (PN 20300021) was applied to each section for 15 min followed by a 10 min wash in Formamide Wash Buffer. Formamide Wash Buffer was removed and replaced with Sample Prep Wash Buffer during MERSCOPE set up. 100 ul of RNAse Inhibitor (New England BioLabs M0314L) was added to 250 ul of Imaging Buffer Activator (PN 203000015) and this mixture was added via the cartridge activation port to a pre-thawed and mixed MERSCOPE Imaging cartridge (VIZGEN PN1040004). 15 ml mineral oil (Millipore-Sigma m5904-6X500ML) was added to the activation port and the MERSCOPE fluidics system was primed according to VIZGEN instructions. The flow chamber was assembled with the hybridized and cleared section coverslip according to VIZGEN specifications and the imaging session was initiated after collection of a 10X mosaic DAPI image and selection of the imaging area. For specimens that passed minimum count threshold, imaging was initiated, and processing completed according to VIZGEN proprietary protocol. Following image processing and segmentation, cells with fewer than 50 transcripts are eliminated, as well as cells with volumes falling outside a range of 100-300um.

The 140 gene Human cortical panel was selected using a combination of manual and algorithmic based strategies requiring a reference single cell/nucleus RNA-seq data set from the same tissue, in this case the human MTG snRNAseq dataset and resulting taxonomy (*1*). First, an initial set of high-confidence marker genes are selected through a combination of literature search and analysis of the reference data. These genes are used as input for a greedy algorithm (detailed below). Second, the reference RNA-seq data set is filtered to only include genes compatible with mFISH. Retained genes need to be 1) long enough to allow probe design (> 960 base pairs); 2) expressed highly enough to be detected (FPKM >= 10), but not so high as to overcrowd the signal of other genes in a cell (FPKM < 500); 3) expressed with low expression in off-target cells (FPKM < 50 in non-neuronal cells); and 4) differentially expressed between cell types (top 500 remaining genes by marker score20). To more evenly sample each cell type, the reference data set is also filtered to include a maximum of 50 cells per cluster.

The main step of gene selection uses a greedy algorithm to iteratively add genes to the initial set. To do this, each cell in the filtered reference data set is mapped to a cell type by taking the Pearson correlation of its expression levels with each cluster median using the initial gene set of size n, and the cluster corresponding to the maximum value is defined as the “mapped cluster”. The “mapping distance” is then defined as the average cluster distance between the mapped cluster and the originally assigned cluster for each cell. In this case a weighted cluster distance, defined as one minus the Pearson correlation between cluster medians calculated across all filtered genes, is used to penalize cases where cells are mapped to very different types, but an unweighted distance, defined as the fraction of cells that do not map to their assigned cluster, could also be used. This mapping step is repeated for every possible n+1 gene set in the filtered reference data set, and the set with minimum cluster distance is retained as the new gene set. These steps are repeated using the new get set (of size n+1) until a gene panel of the desired size is attained. Code for reproducing this gene selection strategy is available as part of the mfishtools R library (https://github.com/AllenInstitute/mfishtools).

H5ad creation: Any genes not matched across both the MERSCOPE gene panel and the mapping taxonomy were filtered from the dataset before starting. From there, cluster means were calculated by dividing the number of cells per cluster by the number of clusters collected. Next, we created a training dataset by finding marker genes for each cluster by calculating the l2norm between all clusters and the mean counts of each gene per cluster. This training dataset was fed into a knn alongside the MERSCOPEs cell by gene panel to iteratively calculate best possible gene matches per cluster. All scripts and data used are available at: https://github.com/AllenInstitute/.

### Analysis

#### Patch-seq data curation and mapping

To evaluate the mapping quality of Patch-seq samples, we calculated the normalized marker sum (NMS) score - a ratio of ‘on’ and ‘off’ markers by subclass (*14*, *16*, *37*). A value of 0.4 was designated as a cut off for low- and high-quality data. Only cells mapping to GABAergic interneuron types and had a NMS score >0.4 were included in the analyses for this study. R

Reference transcriptomic data used in this study were obtained from dissociated inhibitory nuclei collected from human MTG (*1*), and are publicly accessible at the Allen Brain Map data portal (https://portal.brain-map.org/atlases-and-data/rnaseq). This taxonomy consists of a hierarchical dendrogram of cell types, along with a set of marker genes defined to distinguish types at each split in the tree. The Patch-seq transcriptomes were mapped to the reference taxonomy following the ‘tree mapping’ method (map_dend_membership in the scrattch.hicat package) as described previously (*35*, *45*). Briefly, at each branch point of the taxonomy we computed the correlation of the mapped cell’s gene expression with that of the reference cells on each branch, using the markers associated with that branch point (i.e., the genes that best distinguished those groups in the reference), and chose the most correlated branch. The process was repeated until reaching the leaves of the taxonomy To determine the confidence of mapping, we applied 100 bootstrapped iterations at each branch point, and in each iteration 70% of the reference cells and 70% of markers were randomly sampled for mapping. The percentage of times a cell was mapped to a given t-type was defined as the mapping probability, and the highest probability t-type was assigned as the mapped cell type.

#### Morphology feature analysis

Prior to morphological feature analysis, reconstructed neuronal morphologies were expanded in the dimension perpendicular to the cut surface to correct for shrinkage (*69*, *70*) after tissue processing. The amount of shrinkage was calculated by comparing the distance of the soma to the cut surface during recording and after fixation and reconstruction. For mouse cells, a tilt angle correction was also performed based on the estimated difference (via CCF registration) between the slicing angle and the direct pia-white matter direction at the cell’s location (*45*). Features predominantly determined by differences in the z-dimension were not analyzed to minimize technical artifacts due to z-compression of the slice after processing.

Morphological features were calculated as previously described (*45*). In brief, feature definitions were collected from prior studies (*71*, *72*). Features were calculated using the skeleton keys python package (https://github.com/AllenInstitute/skeleton_keys). Features were extracted from neurons aligned in the direction perpendicular to pia and white matter. Laminar axon distribution (bin size of 5 microns) and earth movers distance features require a layer-aligned version of the morphology where node depths are registered to an average interlaminar depth template.

#### Statistical analysis of variability

To assess the variability of morphological features within the acute and culture paradigm, we used a Mann-Whitney U test for each condition. Results were reported as the resulting U statistic and the P-value. P values were corrected for false discovery rate (FDR, Benjamini–Hochberg procedure). Analysis of feature relationships across subclass were assessed using a one-way ANOVA on ranks (KW, Kruskal-Wallis test), correction for FDR and post-hoc Dunn’s tests were run across any pairwise comparison, with an FDR correction when necessary. Analysis of feature relationships with other variables including days in culture and qualitative morpho types were likewise assessed by Mann-Whitney tests for binary variables, KW test for categorical, and Pearson’s correlations for continuous variables, FDR-corrected across features when necessary.

Unless otherwise specified, statistical analyses of morphological features were implemented in python using both the statsmodels and scipy packages. Statistical analyses of electrophysiological features were implemented in Prism or python using both the statsmodels and scipy packages.

### Morphological Clustering

The co-clustering matrix for the SST and PVALB subclass morphology dataset was calculated by iterative random sampling. During each iteration, 95% of samples were randomly selected to create a shared nearest neighbors graph. We then applied the Fast-greedy community detection algorithm using the Python package python-igraph for clustering assignment. For each pair of samples, the co-clustering score was defined as the times of co-clustering normalized by the iterations of co-occurring. Resampling was performed 500 times to reach saturation. Agglomerative clustering using ward linkage was performed on the co-clustering matrix to get clusters.

### Calculating differentially expressed genes

Differentially expressed (DE) genes were calculated for several analyses using the “FindMarkers” function from Seurat V4 (https://satijalab.org/seurat/) (*73*). This function preforms either a Wilcox test or a t-test to calculate a p-value and Bonferroni correction to determine whether genes are DE in two groups. This was used to find DE genes in the following cases: (1) culture microglial genes: cultured cells with and without microglia signature (test.use=“wilcox”, min.pct = 0.5, logfc.threshold = 4); (2) generic microglial genes: common genes with and without microglia signature (wilcox, 0.25, 2) in acute and culture cells; (3) genes increasing or decreasing in PVALB cells in culture: common genes DE in acute vs. culture cells (wilcox, 0.25, 2) with and without microglia signature; and (4) morphology type genes: genes DE between cells with any pair of DBC, MC and Sparse SST morphologies (t-test, 0, 0). Genes DE between dissociated cells in SST and PVALB cell types were calculated using the RNA-seq Data Navigator tool (https://celltypes.brain-map.org/rnaseq/human/mtg), by defining “Set 1 Selection” and “Set 2 Selection” as all PVABL and SST types (excluding SST FRZB) and then running “Find ‘Marker Genes’“.

### Assigning cells a microglial signature

Microglial signature genes were initially defined in cells from culture by following the standard Seurat pipeline for clustering. First, the top variable genes were defined using FindVariableFeatures Seurat function with default parameters. The data was then scaled (using ScaleData), and the dimensionality was reduced by calculating the first 20 principal components (RunPCA), and then generating a 2D UMAP. Finally, cells were clustered into two clusters by running FindNeighbors on the UMAP space and FindClusters with resolution = 0.001. These clusters were used as initial cell sets to define 269 culture microglial genes, as described above. Final microglial calls were defined by rerunning this clustering process independently on cells from culture and cells from acute dissections but using the 269 culture microglial genes as manually input variable genes and setting clustering resolution to 0.1

### Visualization of integrated transcriptomic space

To account for the non-negligible gene expression signatures of culture and microglia, we divided the Patch-seq dataset into four groups (acute, acute - microglial, culture, culture - microglial) and integrated these data together with dissociated reference nuclei by following the Seurat tutorial for integration (*73*). To do this we applied the “FindIntegrationAnchor” function setting the union of all tree mapping marker genes as the set of anchor features, with dissociated cells as the reference data set. We then applied IntegrateData using default parameters to put all cells in the same transcriptomic space. We then scaled the data, reduced the dimensionality using principal component analysis (PCA; 30 PCs), and visualized the results with 2D UMAPs. Finally, metadata such as cluster, paradigm, microglial status, and morphology type were then overlaid onto this UMAP space using different colored or shaped points for all or a subset of cells. Except as noted below, all transcriptomics UMAPs are presented in this UMAP space.

The UMAP in figure S1C was generated using a union of the top 50 generic microglial genes, 50 acute cell markers, and 50 culture cell markers as variable genes, and following the above process without using data integration. Likewise, the UMAP in Figure 3D follows the same procedure only on SST CALB1 cells, setting the 253 genes DE between 10 DBC, MC and Sparse SST cells as variable genes.

---

**Supplementary Figure 1.**
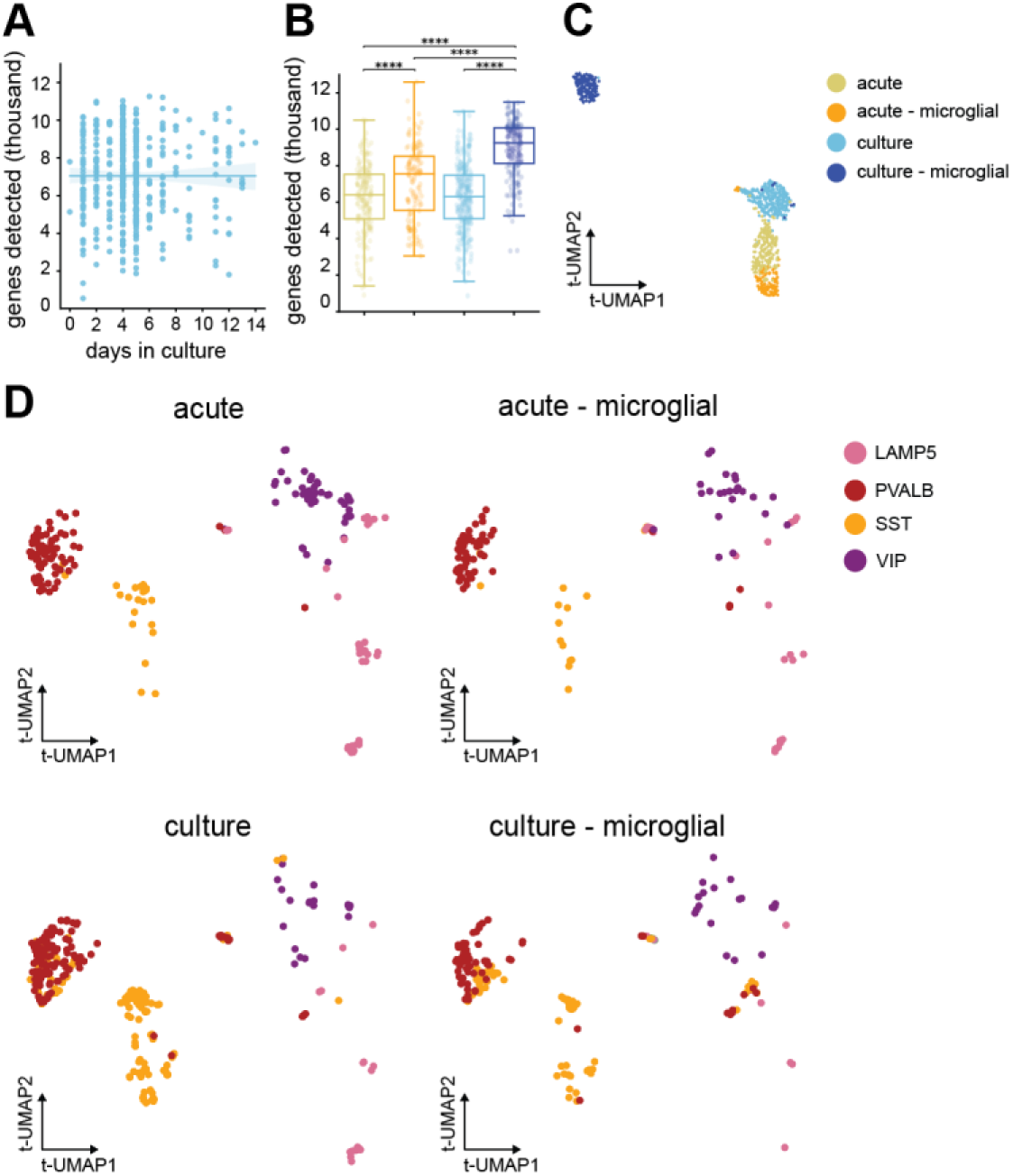
Culture effects and groups expressing microglial markers. (**A**) Scatter plot of number of number of genes versus days in culture for patch-seq experiments (**B**) Box plots represent the difference in the number of genes detected between the two paradigms and microglial groups. (**C**) UMAP representation of transcriptomic space highlighting the paradigm and microglial groups. (**D**) UMAP representation of transcriptomic space separated by paradigm and microglial groups and color-coded by inhibitory subclass.

**Supplementary Figure 2.**
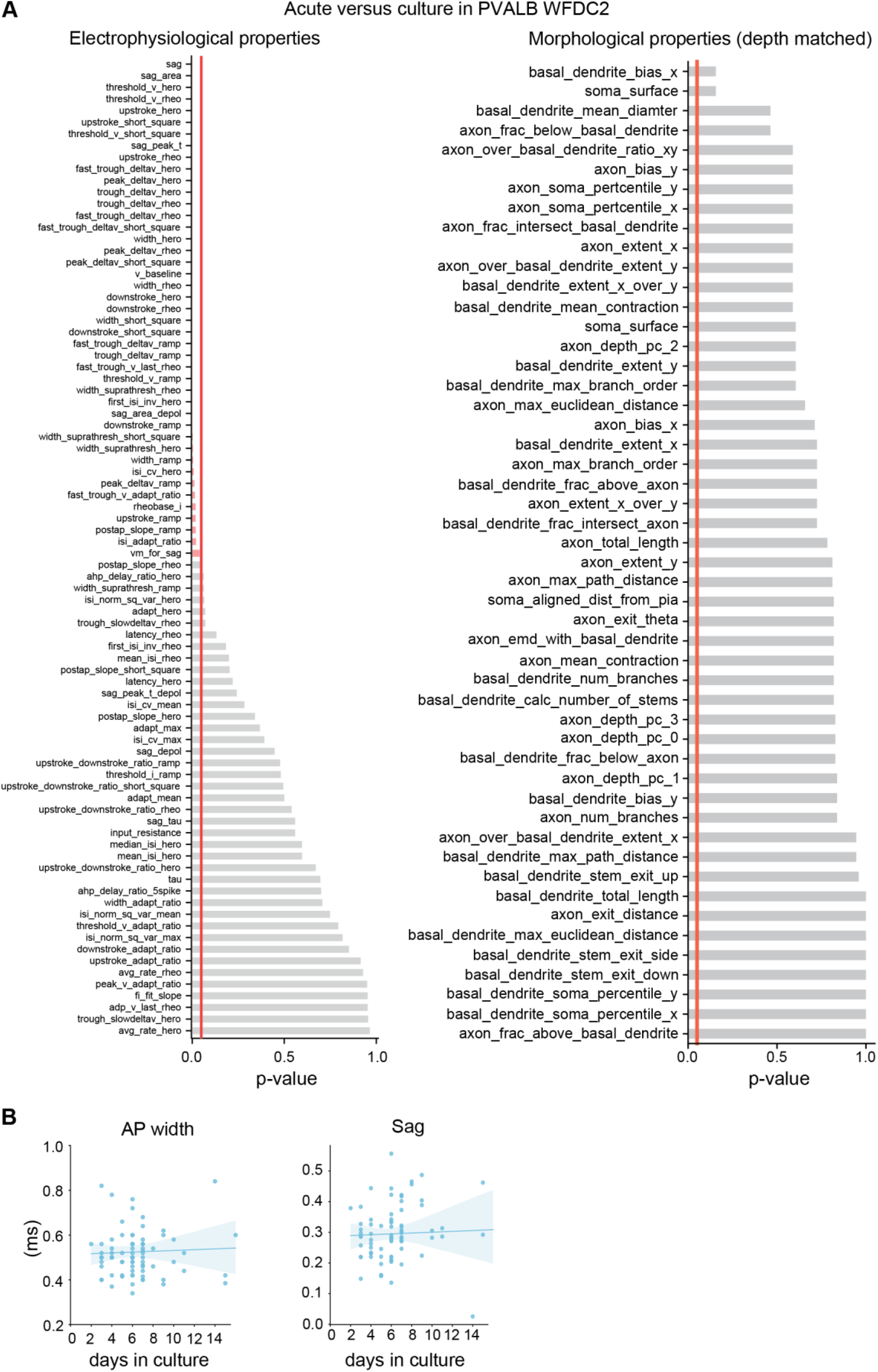
Electrophysiolgoical and morphological changes in PVALB WFDC2 acute and culture groups. **(A)** Mann-Whitney test calculating the level of significance for each morphological feature in acute versus culture for PVALB WFDC2. **(B)** Mann-Whitney test calculating the level of significance for each electrophysiological feature in acute versus culture for PVALB WFDC2. **(C)** Scatter plot of number for sag (left) and action potential width (right) versus days in culture for patch-seq experiments.

**Supplementary Figure 3.**
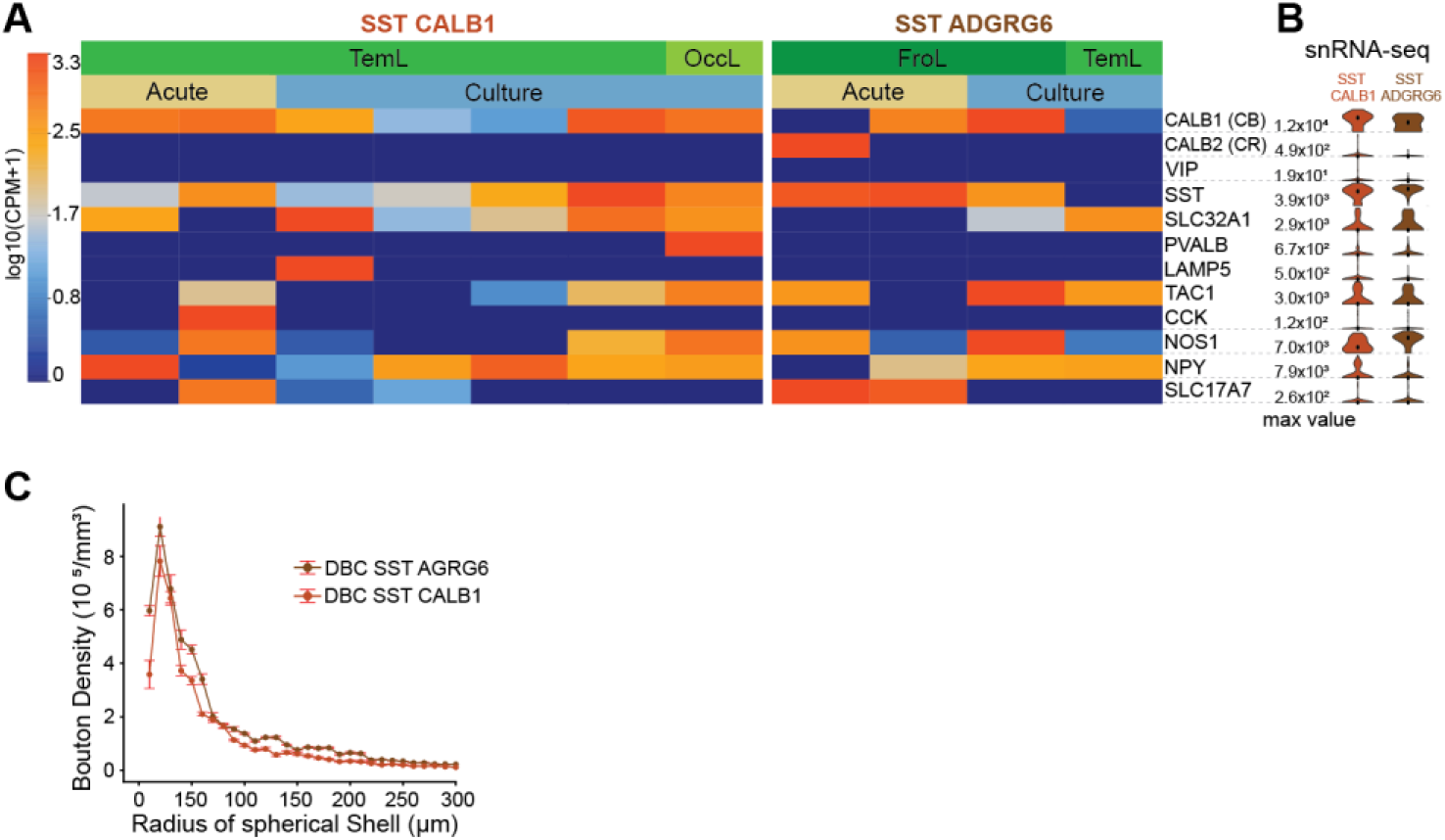
Gene expression in Double bouqet cells and SST CALB1 and SST ADGRG6. **(A)** A heath map dispaying key marker gene expression for Double Bouqet cells at the transcriptomic type level for SST CALB1 and SST ADGRG6, sorted by cortical region and paradigm type. **(B)** Violin plot, at right, shows gene expression for snRNA-seq dataset for the same key marker genes. **(C)** Line plot displaying bouton density of double bouqet cells as a function of radial projection (or distance) per SST CALB1 and SST ADGRG6 types.

**Supplementary Figure 4.**
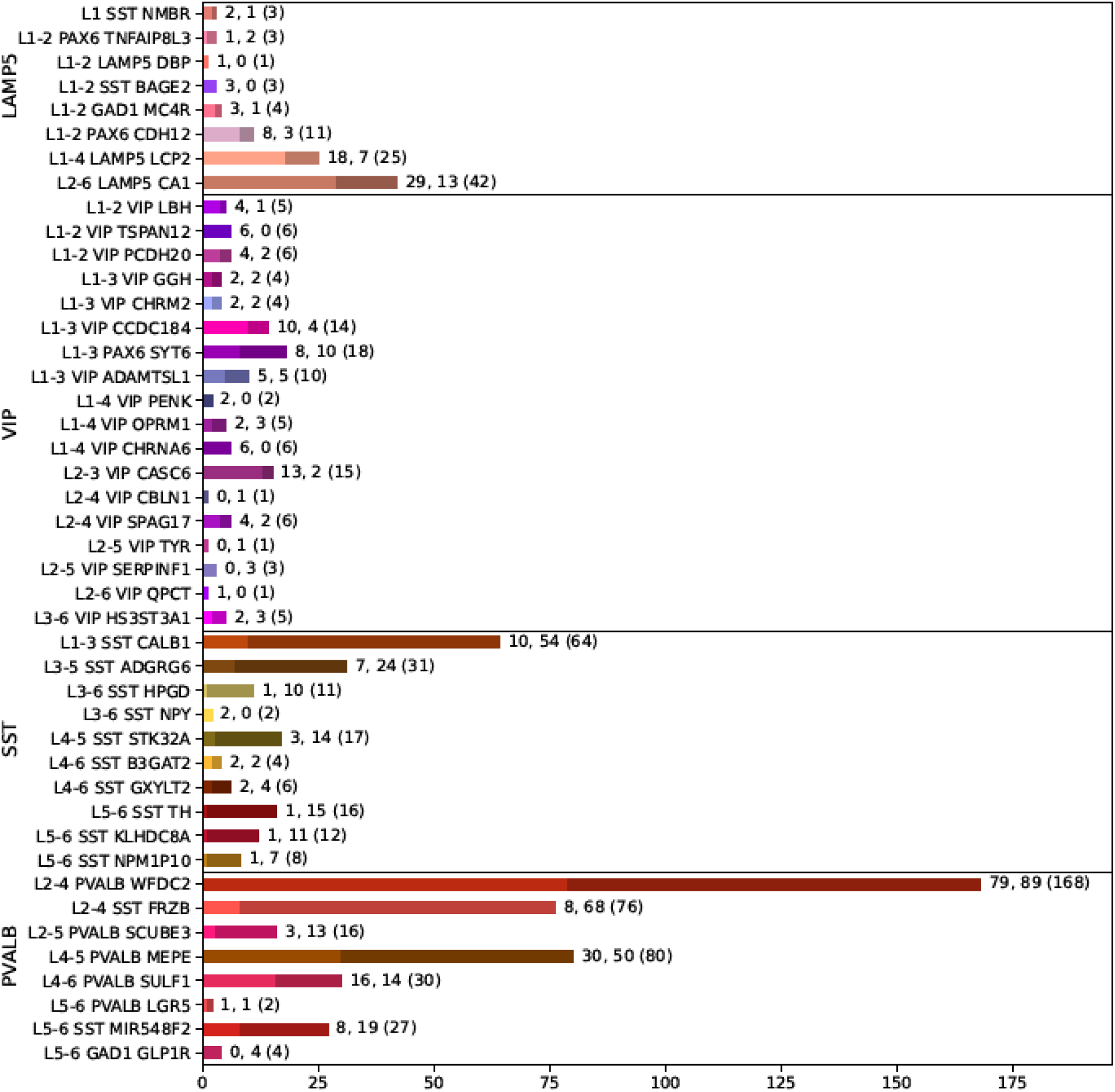
Transcriptomic identify and distribution of manuscript dataset. Patch-seq dataset neurons counts by subclass/t-type and the whether from the acute (first number) or culture (second number) paradigm with total cell count per t-type in parenthesis. Coverage of several transcriptomic types, particular in the SST and PVALB subclasses, was substantially improved through use of the culture paradigm.

**Supplementary Figure 5.**
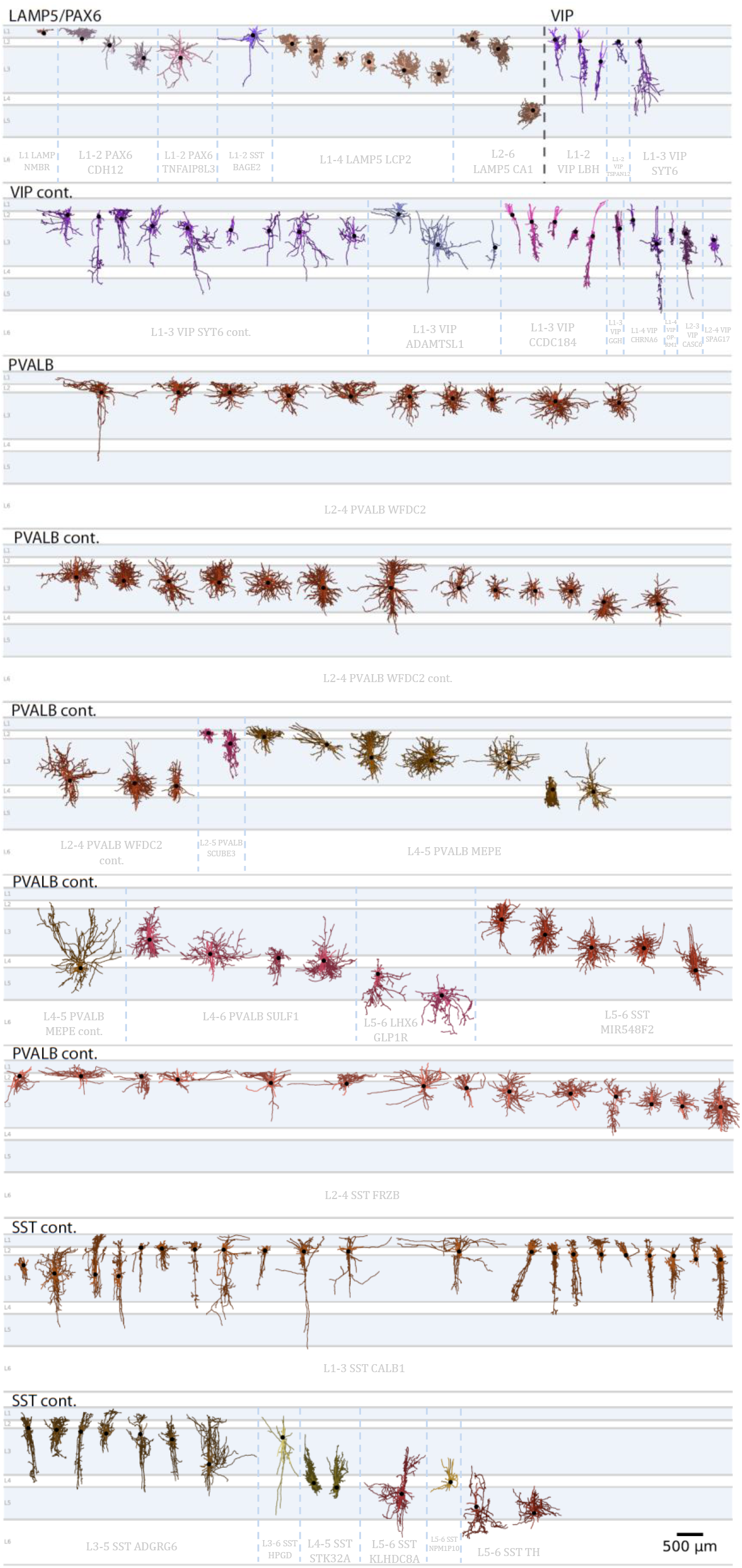
Gallery of morphological reconstructions. All available human dendrite (darker color) and axon (lighter) reconstructions organized by subclass (black text) and t-types (grey text).

